# Immune Stimulation *via* Wounding Alters Chemical Profiles of Adult *Tribolium castaneum*

**DOI:** 10.1101/2021.03.23.436617

**Authors:** Lai Ka Lo, R Reshma, Lisa Johanna Tewes, Barbara Milutinović, Caroline Müller, Joachim Kurtz

## Abstract

Group-living individuals experience immerse risk of disease transmission and parasite infection. Especially in social, but also in some non-social insects, disease control with immunomodulation takes place not only *via* individual immune defenses, but also *via* infochemicals such as contact cues and (defensive) volatiles to mount a group level immunity. However, little is known about whether an activation of the immune system leads to changes in chemical phenotypes, which may mediate these responses. We here asked whether individual immune experience resulting from wounding or injection of heat-killed *Bacillus thuringiensis* leads to changes in the chemical profiles of female and male adult red flour beetles (*Tribolium castaneum*), which are non-social but gregarious. We analyzed insect extracts using GC-FID to study the chemical composition of (1) cuticular hydrocarbons (CHCs) as candidates for the transfer of immunity-related information between individuals *via* contact, and (2) stink gland secretions, with target analysis of benzoquinones as main active compounds regulating ‘external immunity’. Despite a pronounced sexual dimorphism in CHC profiles, wounding stimulation led to similar profile changes in males and females with increases in the proportion of methyl-branched alkanes compared to naïve beetles. While changes in the overall secretion profiles were less pronounced, absolute amounts of benzoquinones were transiently elevated in wounded compared to naïve females. We suggest that the changes in different infochemicals may mediate immune status signaling in the context of both internal and external immune responses in groups of this non-social insect, thus showing parallels to social immunity.

## INTRODUCTION

In group-living animals increased contacts of individuals facilitate the spread of diseases (Kappeler et al. 2015). Thus, the individual ability to respond to immune stimuli and limit infection, but also the recognition of the health-status of group members are vital (Kappeler et al. 2015). Especially eusocial insect species show remarkable physiological and/or behavioral group-level responses to disease, such as distancing (Stockmaier et al. 2021) or hygiene behaviors (Rosengaus et al. 1998; Pull et al. 2018) leading to ‘social immunity’ (Cremer et al. 2018). Insight into individual and group level immunity across insect groups with different types of social structure is highly valuable. However, there have been only a few studies on the phenomenon of social transfer of immunity in non-social insects (Gallagher et al. 2018), and its underlying mechanisms. Horizontal transfer of immunity among conspecifics is often mediated *via* chemicals (Milutinović and Schmitt 2022). Nevertheless, little is known about the direct effects of immune stimuli like wounding and/or pathogen exposure on chemical phenotypes of individuals, which can potentially signal the immune status into the group and/or serve direct defensive functions.

Chemical cues can be crucial for healthy animals to identify diseased conspecifics across taxa (Csata et al. 2017; Pull et al. 2018; Behringer et al. 2020). As one group of such cues acting as contact infochemicals, changes in profiles of cuticular hydrocarbons (CHCs) may play a major role. CHCs cover the surface of insects and consist of complex mixtures of straight-chain, methyl-branched, and unsaturated hydrocarbons (Blomquist and Ginzel 2021). They serve as pheromones or allelochemicals for recognition of species, sex, mutualistic partners, or hosts, and for nestmate recognition and regulation of social interactions in eusocial insects (van Zweden and d’Ettorre 2010; Leonhardt et al. 2016). Many behavioral reactions to immunological and disease cues of nestmates in social insects, and even in a non-social insect species, could be directly related to changes in CHC profiles (Richard et al. 2008; Nielsen and Holman 2012; Baracchi et al. 2012; Hernández López et al. 2017; Alciatore et al. 2021). However, in an ant species immune experiences resulted in changes in the CHCs, but their role in the observed nestmate responses were unclear, suggesting volatile chemicals or behavioral cues as mediators (Alciatore et al. 2021). These findings underline that different infochemicals can be involved in information transfer among conspecifics (Müller et al. 2020).

In addition to contact cues, volatile compounds are known to largely mediate interactions between organisms, both as pheromones detectable over large distances and/or with direct defensive effects against pathogens or predators (Yew and Chung 2015). For example, odors emitted by diseased brood induce hygienic behavior in bees (Swanson et al. 2009), and ant nest volatiles can have antifungal effects (Wang et al. 2015). The release of defensive secretions into the environment can be regarded as ‘external immunity’ (Cotter and Kilner 2010; Otti et al. 2014), i.e., putatively, a process of niche construction (Odling-Smee et al. 2013; Müller et al. 2020). External immunity *via* secretions has not only been found in several social species but also in various group-living (gregarious) insects, e.g. beetles from the Tenebrionidae family (Cotter and Kilner 2010). Their stink gland secretions contain mostly benzoquinones, which, along with further substances, have inhibitory effects on microbes and thus serve a hygienic function (Yezerski et al. 2007; Li et al. 2013). This phenomenon is interpreted as parental care towards larvae (Trivers 1972; Clutton-Brock 1991), but also other adults from the group could benefit from secretions of conspecifics (Gokhale et al. 2017), resulting in herd-protection (Cotter and Kilner 2010). Acting as pheromones, benzoquinones can trigger aggregation in low concentrations, but also mediate dispersal, being repellent and autotoxic in high concentrations (Chapman 1926; Roth and Howland 1941; Sokoloff 1974). However, the cascade from internal immune experience to external immunological responses is rarely studied.

The red flour beetle, *Tribolium castaneum* (Coleoptera: Tenebrionidae), is a non-social, but group-living insect that lives in and consumes flour, and serves as a model species (Campbell et al. 2022), also for studying resistance and immune reactions to various types of stimuli (e.g., Altincicek et al. 2008; Wegner et al. 2008). Exposure to inactivated pathogen or a sublethal dose of living pathogen has previously been shown to induce immune ‘priming’, which is a form of innate immune memory in invertebrates (Little and Kraaijeveld 2004; Netea et al. 2016; Milutinović and Kurtz 2016), leading to enhanced protection (Roth et al. 2009; Tate and Graham 2017; Khan et al. 2017), associated with strong and specific transcriptional responses in *T. castaneum* (Greenwood et al. 2017; Ferro et al. 2019). Beyond that, even after experience of wounding alone this species shows social transfer of immunity, as the interaction of naïve beetles with wounded conspecifics induces an enhanced activity of the important immune effector phenoloxidase (Peuß et al. 2015). However, it is unclear if CHCs or other infochemicals could act as cue for conspecifics to detect immune-challenged beetles. Moreover, *T. castaneum* engages in external immune defenses, releasing benzoquinone-rich stink gland secretions into the flour shared with conspecifics (Prendeville and Stevens 2002; Yezerski et al. 2007; Joop et al. 2014). The role of the main compounds, methyl- and ethyl-1,4-benzoquinone (MBQ and EBQ, respectively) and their carrier 1-pentadecene (Loconti and Roth 1953; Villaverde et al. 2007), for inter- and intraspecific interactions in *Tribolium* beetles is well described (Blum 1981; Yezerski et al. 2007; Verheggen et al. 2007; Duehl et al. 2011). For intraspecific communication, a sex-specific function was suggested for benzoquinones in *T. castaneum*, being specifically involved in female density-dependent competition (Khan et al. 2018). However, to our knowledge it is yet unknown how immune experiences shape overall secretion profiles, whether these responses differ between females and males, and whether potential changes are persistent over time.

We here examined changes in infochemicals of red flour beetles resulting from different immune experiences, i.e., wounding alone or in combination with the injection of heat-killed *Bacillus thuringiensis* bv. *tenebrionis* (bacterial beetle pathogen, ‘priming’ treatment) in comparison to naïve beetles. We analyzed insect extracts using gas chromatography flame ionization detection and expected changes in (1) the CHC profiles, which may act as a cue for the immune status of conspecifics and (2) the gland secretion profiles, in particular quantities of EBQ, MBQ and 1-pentadecene, which are involved in external immunity and population regulation.

## METHODS AND MATERIALS

### Model Organisms

Our study was done on the outbred strain Croatia 1 (CRO1) of *T. castaneum* which was kept at 30 °C, 70% humidity and a 12 h:12 h light:dark cycle in plastic boxes containing heat-sterilized (75 °C for at least 24 hours) organic wheat flour (Bio Weizenmehl Type 550, dm-drogerie markt) with 5% brewer’s yeast powder (hereafter the flour-yeast mixture is referred to as “flour”) (Milutinović et al. 2013). For this experiment, around 2,000 one-month old adult beetles were allowed to mate and oviposit for 24 h in approximately 350 g flour.

A culture of *Bacillus thuringiensis* bv. *tenebrionis* was obtained from Bacillus Genetic Stock Center (Ohio State University, USA). The species is a bacterial pathogen of beetles, and immune priming *via* injection of this bacterium has been demonstrated in previous work (e.g., Ferro et al. 2019). Heat-killed vegetative cells at 1 × 10^9^ cells mL^-1^ used for the immune stimulation treatment were prepared as previously described (Roth et al. 2009; Ferro et al. 2017) (see Methods S1 for details).

### Immune Treatment

Ten days after oviposition, all experimental beetles were individually placed into sterile 96-well plates (Type F, Sarstedt) containing 0.08 g flour per well to minimize any injuries due to conspecific cannibalism or mating (Park et al. 1965; Via 1999; Alabi et al. 2008). One week post eclosion, virgin beetles of both sexes were divided into three treatment groups: Untreated as the handling control (‘naïve’), injected with sterile PBS (‘wounded’) and heat-killed *B. thuringiensis* bv. *tenebrionis* injected (‘primed’). For the latter two treatments, 18.4 nL (~ 20,000 cells per injection in the ‘primed’ treatment) of solution was injected into the body cavity of adults with a nanoinjector (Drummond Nanoject II) between the head and pronotum laterally towards the abdomen. Subsequently, beetles were transferred into clean 96-well plates without flour, to reduce the contamination of samples by faecal material and flour.

For the profiling of CHCs at 18 h post treatment, we prepared n = 6 samples per sex and treatment with four beetles pooled into each sample. The sampling time was chosen based on a previous study that reported transfer of immunity within 18 h post treatment (Peuß et al. 2015). For the stink gland secretion collection at 24 h and 72 h post treatment, we obtained n = 8 samples of individual beetles per sex, treatment and time point. These two time-points were chosen to test the time course of the stink gland responses to the treatment.

### Extraction of Beetle Chemicals

For extraction of the CHCs, four beetles per sample were pooled and freeze-killed at −20 °C. Samples were shaken in 120 μL of *n*-hexane (GC-MS grade, Merck) containing 5 μL of eicosane (99.5%, Sigma-Aldrich) solution (0.2 mg mL^-1^ in *n*-hexane) as internal standard at 4 °C for 10 min. After centrifugation, 100 μL of each extract was dried under reduced pressure and resolved in 80 μL of *n*-hexane. The volume of *n*-hexane was scaled down accordingly in three samples, where less than four beetles were available. For each set of samples (secretions and CHCs), four blank samples were prepared alongside with the beetle samples but did not contain beetles. For extraction of stink gland secretions, individual beetles were transferred to tubes that were immersed in ice water at 3 °C for 3 min to stimulate the release of secretions (Joop et al. 2014). Beetles were then freeze-killed at −20 °C and shaken in 65 μL ice-cold acetone (LC-MS grade, Sigma-Aldrich) containing 0.05 mg mL^-1^ octadecane (98.5%, Sigma-Aldrich) as internal standard at 4 °C for 5 min. After centrifugation, 50 μL of each extract were kept at −20 °C until analysis.

### Chemical Analysis, Data Pre-processing and Feature Identification

All samples were analyzed using a gas-chromatograph coupled with a flame ionization detector (GC-FID-2010 Plus, Shimadzu) and equipped with a VF-5ms column (30 m × 0.25 mm × 0.25 μm, with 10 m EZ-guard column, J&W Agilent Technologies). For the analysis of CHCs, 1 μL from each sample was injected with a split ratio of 2 at a constant nitrogen flow of 1.02 mL min^-1^. The column oven temperature started at 100 °C, first increased to 230 °C with a rate of 20 °C min^-1^, then increased to 260 °C with a rate of 5 °C min^-1^, further increased to 277 °C with a rate of 1 °C min^-1^, and finally increased to 320 °C with a rate of 5 °C min^-1^, which was held for 9.9 min. For the analysis of stink gland secretions, a sample volume of 1 μL was injected with a split ratio of 3 at a constant stream of hydrogen flow of 1.13 mL min^-1^. The column oven temperature was set at 50 °C, held for 2 minutes and increased to 300 °C with a rate of 10 °C min^-1^, followed by a holding time of 10 min. Additionally, two alkane mix standards, C8-20 and C8-C40 (both Sigma-Aldrich), were analyzed using both chromatographic methods. Standards of MBQ (98% purity), methyl-1,4-hydroquinone (99%, both Sigma-Aldrich) and ethyl-1,4-hydroquinone (97%, Abcr) were analyzed using the method for gland secretions. The hydroquinones are the form to which benzoquinones convert to (Yezerski et al. 2000) and could thus appear in the datasets.

Data pre-processing was done using the GCsolution Postrun software (Version 2.30.00, Shimadzu), and retention time alignment of peaks in R (version 4.0.3, R Core Team 2020) using the package *GCalignR* (Ottensmann et al. 2018); for feature selection criteria and details on settings see Methods S2. Data were normalized to the internal standard and relative feature values calculated (proportion of total amount per sample). For identification, the retention times of peaks from the alkane mix standards were used to calculate the retention index (RI) of detected features according to (Kováts 1958). Putative identification of CHCs was done by comparing the RIs with those published in previous studies (Lockey 1978; Alnajim et al. 2019; Awater-Salendo et al. 2020). Benzoquinones (BQs) as well as other major secretion compounds previously described in *T. castaneum* glandular secretions were identified by comparing their RIs to previous studies (Li et al. 2013; Lehmann 2015) and by comparisons to the analyzed commercial standards.

### Statistical Analyses

All statistical analyses were done using R (version 4.0.3, R Core Team 2020). To visualize dissimilarities between groups of beetles, non-metric multidimensional scaling (NMDS) based on pairwise Kulczynski distances (package *vegan*, version 2.5-7, Oksanen et al. 2020), was applied to the relative datasets (Wisconsin-square-root-double-standardized) of CHCs and secretions, respectively. After testing assumptions, permutational multivariate analysis of variance (PERMANOVA) was performed to test for the influence of treatment on the dissimilarities, partly combined with post hoc comparisons (Martinez Arbizu 2020) including corrections (Benjamini and Hochberg 1995) (see Methods S3). Analyses of treatment effects were done separately for sexes and in secretion datasets also separately for time points. Additionally, sex differences in the CHC profiles of naïve females *versus* naïve males were addressed using PERMANOVA.

Responses of CHC profiles of females and males to treatments were visualized in heatmaps using the package *pheatmap* (Kolde 2019). Therefore, log_2_ fold change values were calculated for all CHCs and replicates in comparison to the median value of the naïve female or male beetles and data plotted as heatmaps. Moreover, heatmaps comparing CHC profiles of naïve females and naïve males were generated based on z-scores for each CHC. All heatmap plots included Euclidean cluster analysis.

For the secretions, we additionally identified features most important for profile differences between treatment groups within females and males 24 h after treatment using a conditional random forest classification (for details see Methods S3) (Strobl et al. 2009a, b; De Moraes et al. 2014). The relative amounts of these features, as well as the absolute amounts of MBQ, EBQ and 1-pentadecene (1-C15-ene), were analyzed for an influence of treatment within females and males at both time points. We tested the model assumptions (see Methods S3), and either fitted linear models with appropriate transformations revealed using the package *bestNormalize* (Peterson 2021), or performed non-parametric Kruskal-Wallis tests. When significant treatment effects were founds, Benjamini-Hochberg-corrected post hoc comparisons were performed between treatment groups.

## RESULTS

### Cuticular Hydrocarbons

In total, 20 CHCs were putatively identified in the dataset. These consisted of seven *n*-alkanes (C25-C31), six 3- and 4-methyl-branched alkanes and seven internally methyl-branched alkanes of which one was a dimethyl alkane (Table S1). Significant immune treatment effects on the CHC profile were found for both sexes (Table 1). In both females and males, the CHC profiles of naïve beetles differed from those of immune treated beetles, but separated stronger from profiles of wounded than of primed beetles (Fig. 1a,c). Accordingly, in post hoc tests on datasets of both females and males, CHC profiles of naïve beetles differed significantly from those of wounded, but not from those of primed beetles, while CHC profiles of wounded and primed beetles were quite similar (Table 1). Interestingly, despite general sex differences (Fig. S1), the same CHCs were overall involved in separation of naïve from wounded and primed beetles in females and males (Fig. 1). As inferred from the CHC scores in the NMDS plot and heatmaps showing treatment-related differences relative to naïve beetles, higher proportions of shorter methyl alkanes (C26-C29) characterized CHC profiles of wounded and primed beetles in comparison to those of naïve beetles (Fig. 1). The CHC profiles of naïve beetles had higher proportions of comparatively longer *n*-alkanes (C29-C31) and also of 3-MeC31 (Fig. 1).

**Table 1.**
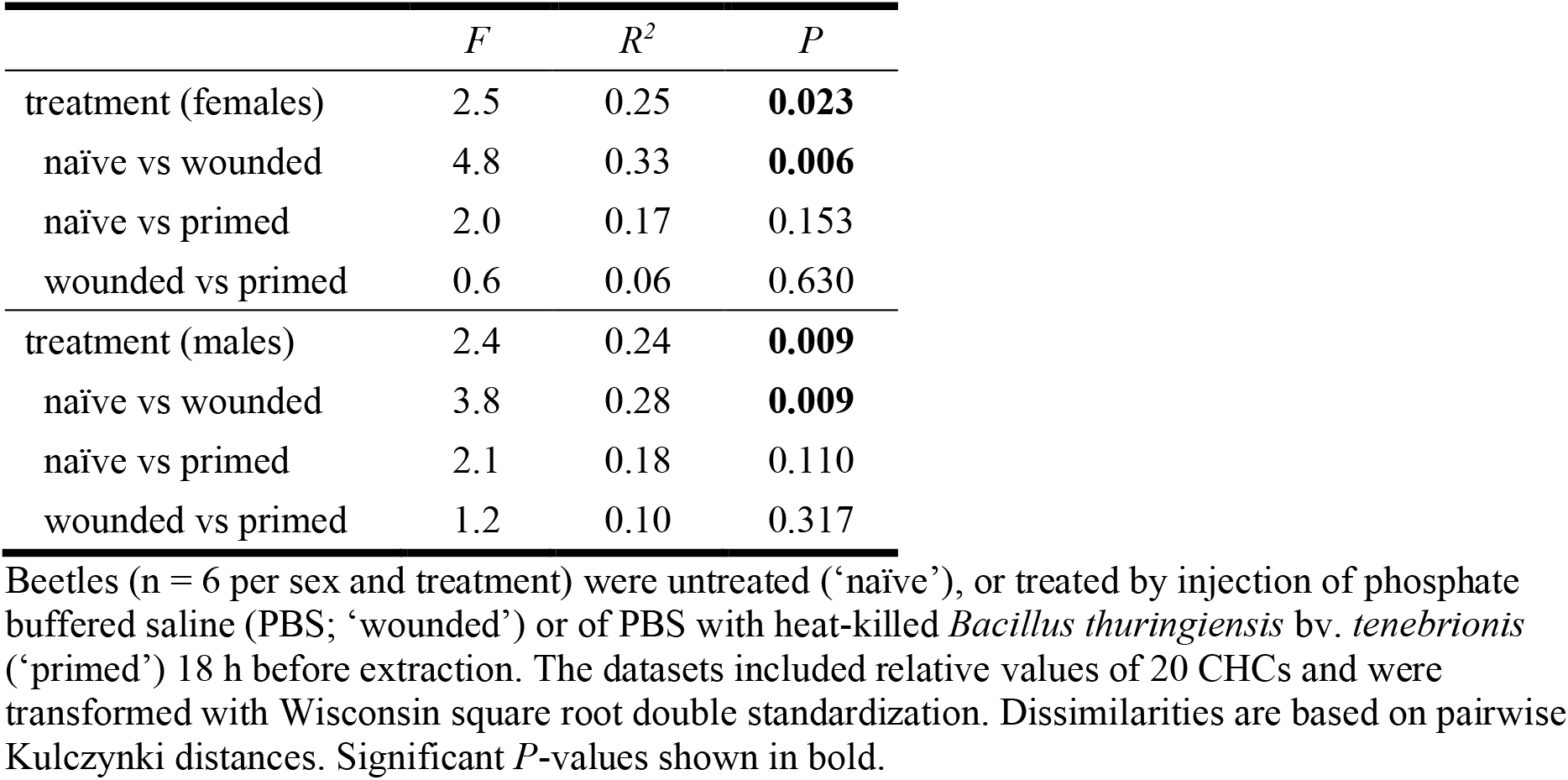
Effects of treatment on the composition of CHC profiles in female and male *Tribolium castaneum* beetles in a permutational analysis of variance (overall tests and pairwise comparisons)

|  | *F* | *R*<sup>2</sup> | *P* |
| --- | --- | --- | --- |
| treatment (females) | 2.5 | 0.25 | **0.023** |
| naïve vs wounded | 4.8 | 0.33 | **0.006** |
| naïve vs primed | 2.0 | 0.17 | 0.153 |
| wounded vs primed | 0.6 | 0.06 | 0.630 |
| treatment (males) | 2.4 | 0.24 | **0.009** |
| naïve vs wounded | 3.8 | 0.28 | **0.009** |
| naïve vs primed | 2.1 | 0.18 | 0.110 |
| wounded vs primed | 1.2 | 0.10 | 0.317 |

Beetles (n = 6 per sex and treatment) were untreated ('naïve'), or treated by injection of phosphate buffered saline (PBS; 'wounded') or of PBS with heat-killed *Bacillus thuringiensis* bv. *tenebrionis* ('primed') 18 h before extraction. The datasets included relative values of 20 CHCs and were transformed with Wisconsin square root double standardization. Dissimilarities are based on pairwise Kulczynski distances. Significant *P*-values shown in bold.

**Fig. 1.**
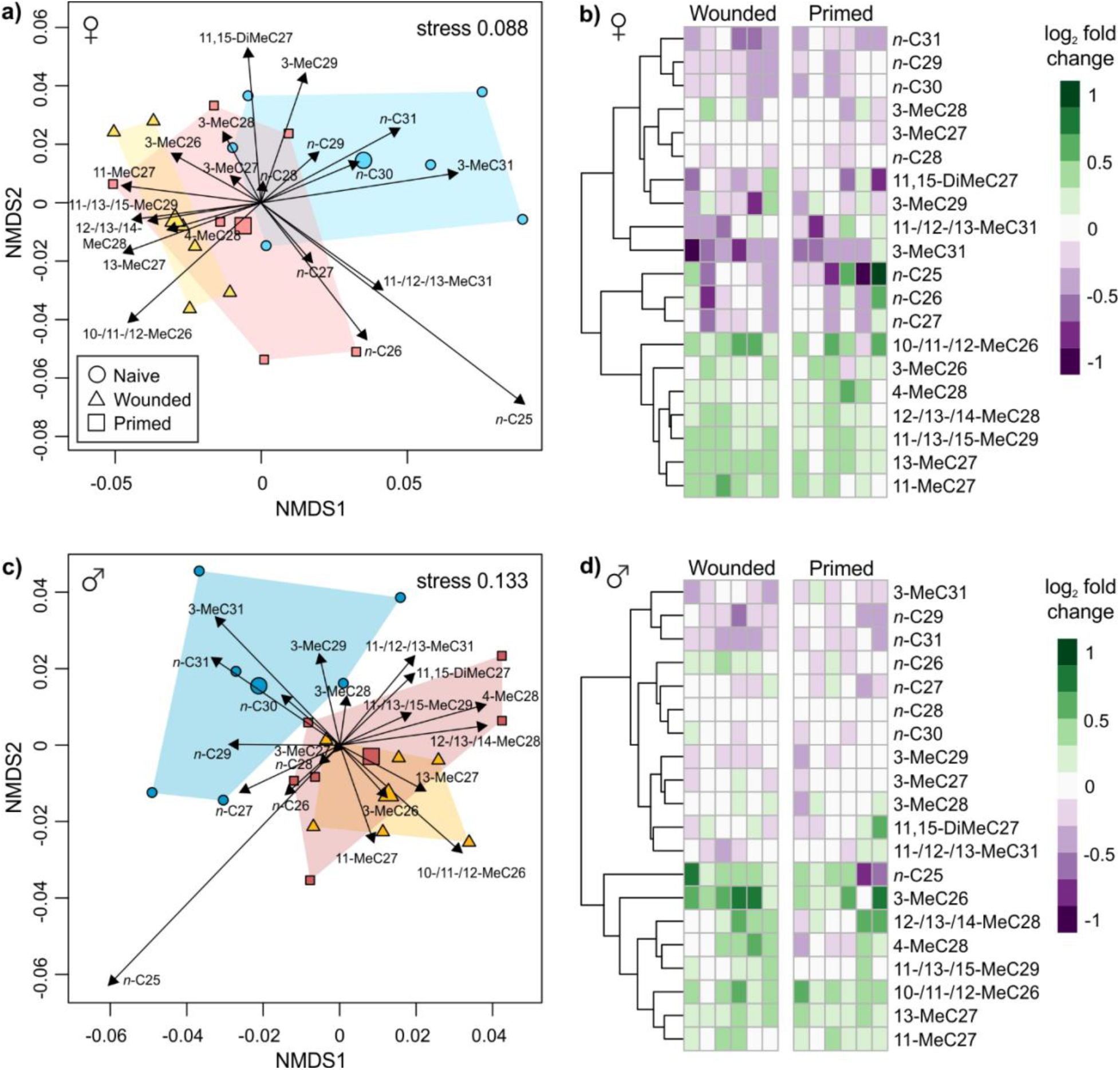
NMDS plots (a,c) and heatmaps (b, d) of cuticular hydrocarbon (CHC) profiles from female (**a,b**) and male (**c,d**) beetles of *Tribolium castaneum*. Beetles (n = 6 per sex and treatment) were untreated (‘Naïve’), or treated by injection of phosphate buffered saline (PBS; ‘Wounded’) or of PBS with heat-killed *Bacillus thuringiensis* bv. *tenebrionis* (‘Primed’) 18 h before extraction. All plots are based on relative amounts of 20 individual CHCs. For NMDS plots, datasets were transformed with Wisconsin square root double standardization and dissimilarities are based on pairwise Kulczynki distances. Data points represent profiles of individual beetles, larger symbols show centroids for treatment groups. Arrows range from zero to positions of expanded weighted average scores of CHCs. Heatmaps show increase and decrease in the proportion of individual CHCs in wounded and primed females (**b**) and males (**d**) in relation to the median value of female or male naïve beetles as log_2_ fold change. Clustering of CHCs is based on Euclidean distances

NMDS plots of CHC profiles of naïve females *versus* males resulted in a clear separation of the sexes (Fig. S1a), which was mirrored in a significant difference (PERMANOVA; *F* = 13.5, *R^2^* = 0.58, *P* = 0.002). Most obviously, the CHC profiles of female beetles were characterized by higher proportions of rather short-chain methyl alkanes, especially of the cluster comprising 10-/11-/12-MeC26, 3-MeC27 and 13-MeC27 (Fig. S1a,b). In contrast, male CHC profiles harbored higher proportions of rather long-chain *n*- and methyl alkanes, especially of the cluster comprising 3-MeC29, *n*-C30 and *n*-C31 (Fig. S1a,b).

### Stink Gland Secretions

For analyses of the dataset acquired 24 h after treatment, 26 chemical features were included, of which 24 were also considered in the dataset acquired 72 h after treatment (Table S2). The benzoquinones MBQ and EBQ could clearly be identified, but the corresponding hydroquinones were not detected in the dataset. The other five identified features were alkenes with one or two double bonds (Table S2). The amounts of benzoquinones, together with 1-C15-ene made up the major part of the overall secretion composition (Table S2).

For secretion profiles of female beetles 24 h after treatment, the treatment effect was marginally non-significant (PERMANOVA, *F* = 2.0, *R^2^* = 0.17, *P* = 0.057). As visible in the NMDS plot, this tendency resulted mainly from separation of naïve and wounded females (Figs 2a, S2). Analyzing male beetles sampled 24 h after treatment, the composition of the overall secretions seemed visually quite similar in all three treatment groups (Figs 2c, S3), and there was no significant treatment effect (*F* = 1.4, *R^2^* = 0.12, *P* = 0.180). To explore if individual features from the profiles respond to immune challenge in females *versus* males, random forest analysis was applied for both sexes separately. Random forest analysis of female beetles revealed one unidentified feature, RI 1015, as important for separation of treatment groups (Fig. S4a). The relative amount of this feature was significantly influenced by treatment in females (*Kruskal-Wallis test*; 2 df, *χ^2^* = 9.84, *P* = 0.034; males: 2 df, *χ^2^* = 1.72, *P* = 0.490) and lower in both primed and wounded beetles compared to naïve ones (Fig. S4b). The unidentified feature RI 1473 was suggested as important in males (Fig. S5a). The relative amount of this feature was significantly influenced by treatment in males (*Kruskal-Wallis test*; 2 df, *χ^2^* = 9.27, *P* = 0.034; females: 2 df, *χ^2^* = 3.43, *P* = 0.370), being higher in wounded than in naïve beetles (Fig. S5b). In the dataset acquired 72 h after treatment, neither females (PERMANOVA, *F* = 0.6, *R^2^* = 0.105, *P* = 0.802) nor males (*F* = 0.6, *R^2^* = 0.06, *P* = 0.854) revealed statistical or visual effects of treatments on the secretion profiles (Fig. 2b,d) and thus further random forest analysis was not performed.

**Fig. 2.**
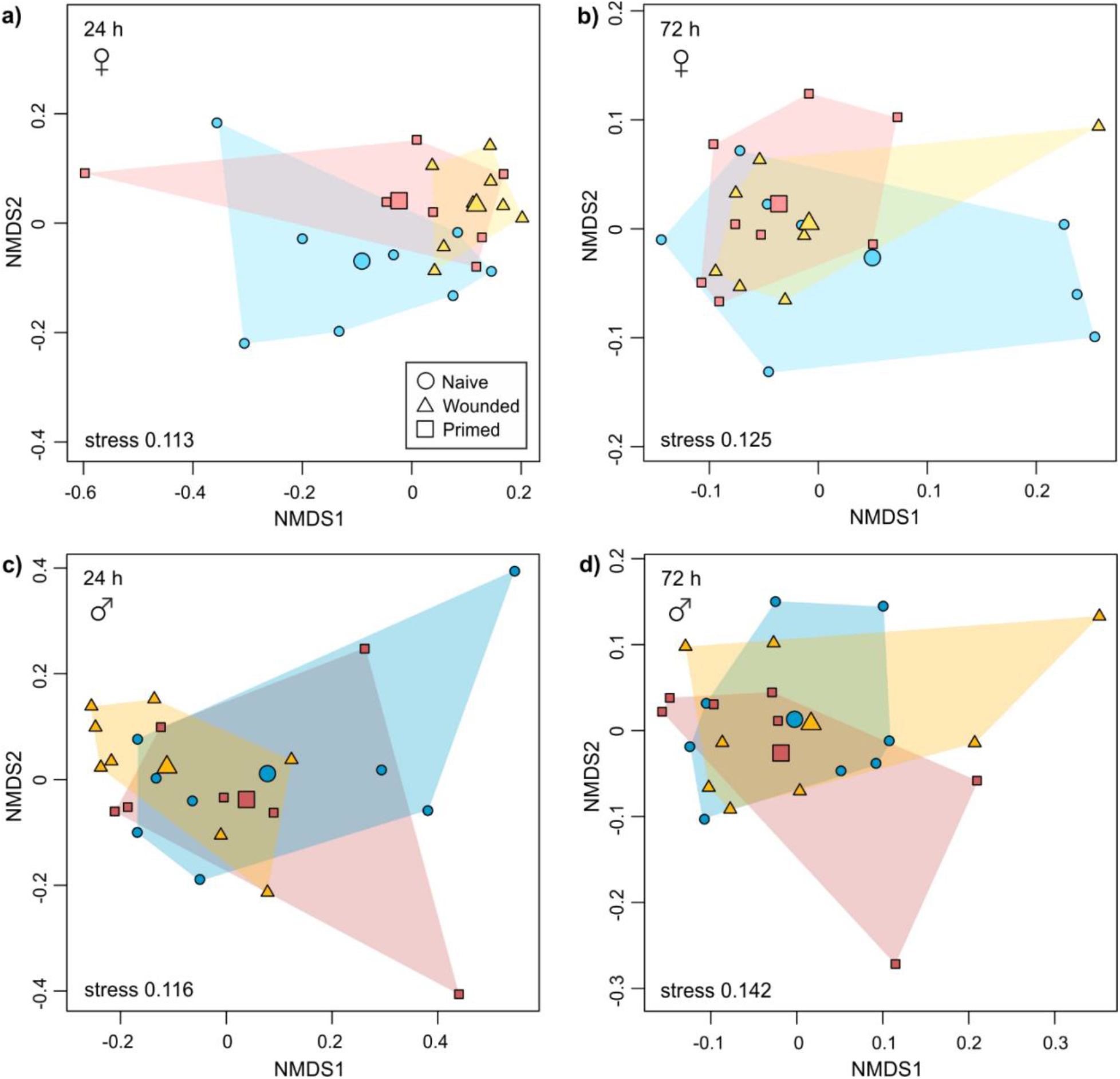
NMDS plots of glandular secretion profiles from female (**a**, **b**) and male (**c**, **d**) beetles of *Tribolium castaneum* of different immune treatment groups. Beetles (n = 7-8 per sex and treatment) were untreated (‘Naïve’), or treated by injection of phosphate buffered saline (PBS; ‘Wounded’) or of PBS with heat-killed *Bacillus thuringiensis* bv. *tenebrionis* (‘Primed’) 24 h (**a**, **c**) or 72 h (**b**, **d**) before extraction. Plots are based on relative amounts of 26 individual chemical features released from glands. Datasets were transformed with Wisconsin square root double standardization. Dissimilarities are based on pairwise Kulczynki distances. Data points represent profiles of individual beetles, larger symbols show centroids for treatment groups

Furthermore, we compared the absolute amounts of the major compounds MBQ, EBQ and their carrier 1-C15-ene. In the secretions acquired after 24 h, an effect of treatment was only significant for females, but not for males in these three compounds (Table 2, Fig. 3). In post hoc tests, the absolute amounts of all three compounds differed significantly between naïve and wounded females (Table 2), being higher in wounded individuals (Fig. 3). The amounts of MBQ and EBQ were by trend also higher in wounded than in primed females and this difference was significant for 1-C15-ene (Table 2, Fig. 3). In the dataset acquired 72 h after treatment, no treatment effects were found in females or males (Table 2). Interestingly, in secretions collected after 72 h the benzoquinones in primed, not in wounded females reached the highest average amounts (Fig. 3).

**Table 2.**
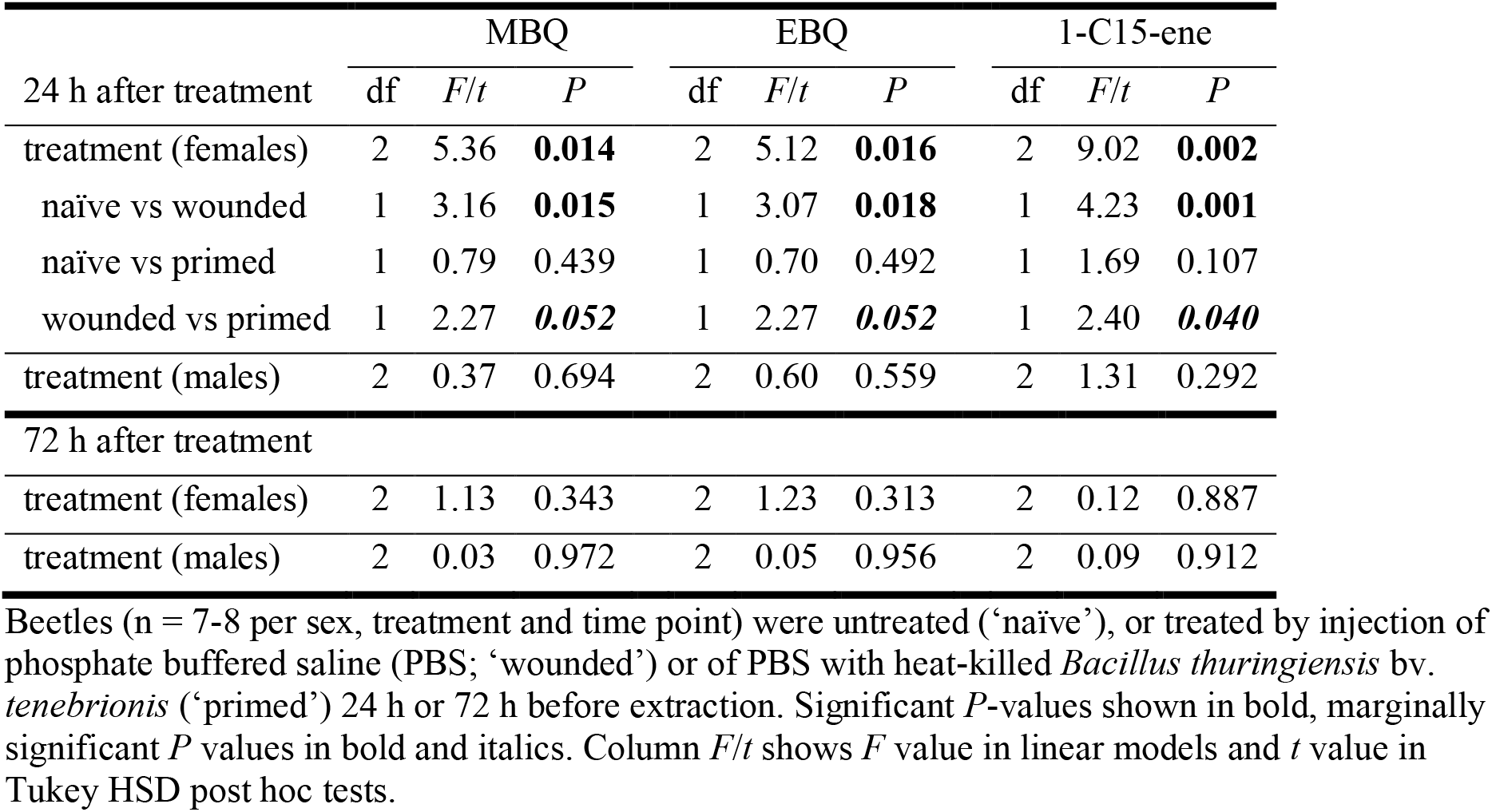
Effects of treatment on absolute amounts of methyl-1,4-benzoquinone (MBQ), ethyl-1,4-benzoquinone (EBQ), and 1-pentadecene (1-C15-ene) from glandular secretions of female and male beetles of *Tribolium castaneum* in linear models (overall tests) and Tukey HSD post hoc tests (pairwise comparisons)

**Fig. 3.**
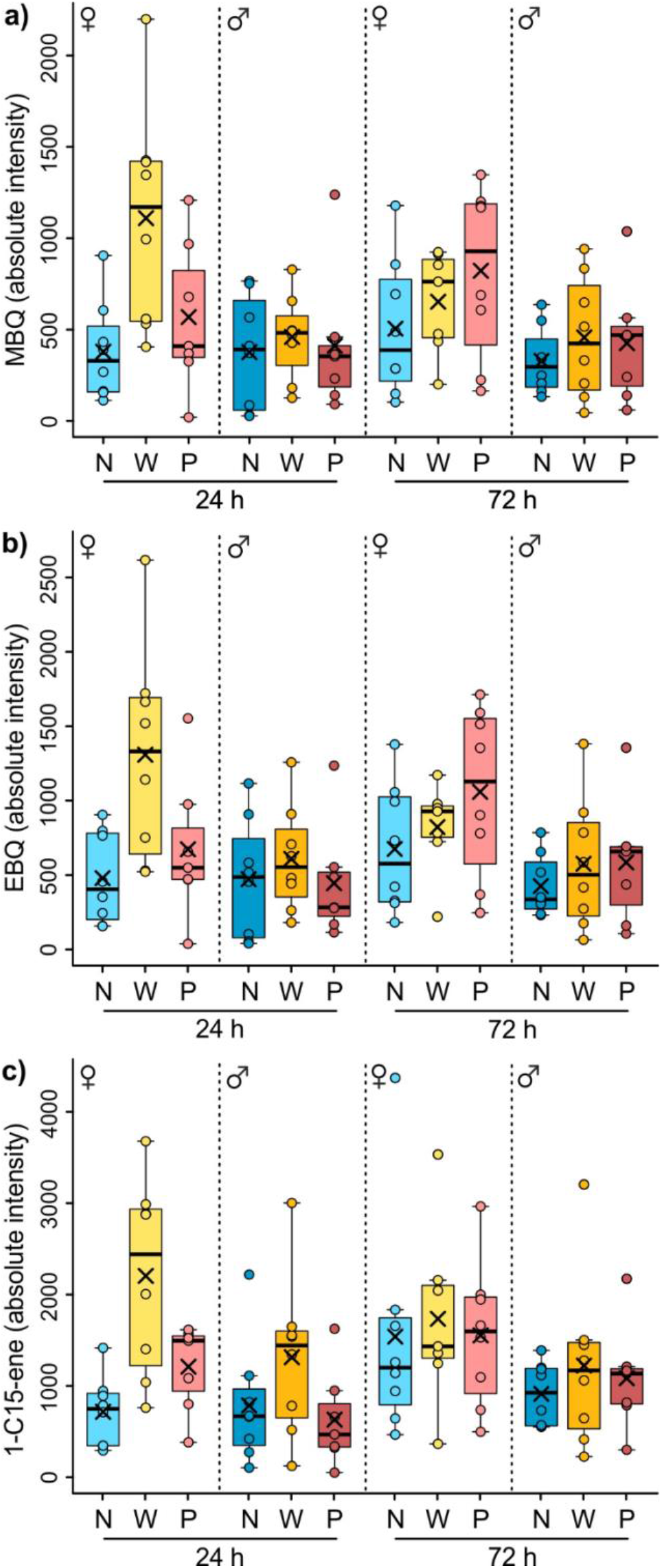
Amount of methyl-1,4-benzoquinone (MBQ), ethyl-1,4-benzoquinone (EBQ) and 1-pentadecene (1-C15-ene) from glandular secretions of female and male beetles of *Tribolium castaneum* at two time points (24 h, 72 h) after treatment. Beetles (n = 7-8 per sex, treatment and time point) were untreated (naïve, ‘N’), or treated by injection of phosphate buffered saline (PBS; wounded, ‘W’) or of PBS with heat-killed *Bacillus thuringiensis* bv. *tenebrionis* (primed, ‘P’) 24 h or 72 h before extraction. Boxes show 25th and 75th percentiles, lines are medians, crosses are means, whiskers mark minimum and maximum within 1.5-fold interquartile ranges. Dots represent values of individual beetles

## DISCUSSION

As group-living animals that share the same food resources and live in proximity, flour beetles are prone to increased disease transmission. We investigated the influence of different immune stimuli on profiles of different infochemicals in *T. castaneum*. Wounding led to similar changes in CHC patterns of females and males, potentially involved in transferring information on the immune status within the group. Wounding-induced changes in gland secretion compounds that could directly mediate external immunity and behavioral responses were mainly restricted to females. The response to priming was lower than to wounding across both types of infochemicals, emphasizing the complexity of physiological responses, especially regarding internal (individual) *versus* external (group) immunity.

The CHC profiles of the beetles were hypothesized to be altered upon immune stimulation *via* wounding and/or priming. Indeed, we found significant differences in the CHC profiles of naïve *versus* wounded females and males. These changed profiles may act as a potential cue for the detection of immune experience in conspecifics, leading to the previously described social transfer of immunity (enhanced phenoloxidase activity) from wounded to naïve *T. castaneum* during cohabitation (Peuß et al. 2015). Moreover, in that study cohabitation with wounded beetles resulted in reduction in expression of two heat shock protein genes, *Hsp83* and *Hsp90*, in naïve conspecifics (Peuß et al. 2015) within the same time span, in which profile changes occurred in the present study. As the molecular chaperone HSP90 serves as an evolutionary capacitor by mediating the storage and release of cryptic genetic variation (Rutherford and Lindquist 1998), such down-regulation has the potential to lead to increased adaptability. However, a causal relationship between changes in CHCs and transfer of information on wounding, including the consequences for social immunity and evolutionary capacitance, needs to be proven in additional bioassays.

The CHC profiles of immune-stimulated beetles in the present study, especially the wounded ones, were characterized by higher proportions of shorter methyl-branched alkanes. This is in line with studies on other insect species, in which specifically methyl alkane CHCs were altered by immune challenge (Richard et al. 2008; Baracchi et al. 2012; Alciatore et al. 2021), and could therefore contribute to social transfer of immunity. Despite higher metabolic costs for these CHCs (Nelson 1993), a generally more important role of methyl alkanes in comparison to linear *n*-alkanes as interspecific recognition cues is well documented in social insects (van Zweden and d’Ettorre 2010). In larvae of *T. castaneum*, wounding caused an upregulation of multiple candidate fatty acid synthase genes (Behrens et al. 2014), which are involved in the synthesis of methyl alkanes (Blomquist and Ginzel 2021; Holze et al. 2021). Assuming that CHC synthesis is regulated in comparable ways in *T. castaneum* larvae and adults, larvae may carry equivalent information on the immune status in their CHCs profiles. Moreover, the wounding-induced alterations in methyl alkanes in adult CHCs observed in the present study may then result from similar, active processes as suggested for the larvae. Methyl alkanes have furthermore been shown to directly mediate interspecific interactions in larvae of *T. castaneum*, acting as host detection cues for parasitoids (Awater-Salendo et al. 2020). Taken together, methyl-branched CHCs are very likely an important factor also in intraspecific transfer of information in both larvae and adults of *T. castaneum*.

Furthermore, we could demonstrate a sexual dimorphism in the CHC profiles of naïve *T. castaneum* adults, which is also well documented in various other invertebrate species (Jallon and David 1987; Thomas and Simmons 2008; Müller and Müller 2016). However, CHC profiles seemed to change in similar directions in *T. castaneum*. In contrast, in the non-social species *T. molitor* changes of the CHC profiles after immune challenge were sex-specific, and the resulting (sexual) signals towards conspecifics resulted in advantages for the wounded males (Nielsen and Holman 2012), interpreted as terminal investment (Sadd et al. 2006; Nielsen and Holman 2012; Gallagher et al. 2018). The similarity in the CHC profile changes across sexes in the present study, potentially equivalently displaying wounding experience, may be beneficial for an effective transfer of information into the group, and could thus be regarded as ‘social trait’. To investigate the potential of CHCs to signal the immune status to conspecifics and their species-specific ecological functions, studies are needed in both sexes of various eusocial *versus* non-social insects.

Flour beetles perform niche construction by secreting quinones-rich stink gland secretions into the flour, which influences microbial growth (Van Wyk et al. 1959; Sokoloff 1974) as a form of external immunity (Joop et al. 2014; Otti et al. 2014). Considering this as parental care (Trivers 1972; Clutton-Brock 1991), we expected that adult females and/or males may modulate their secretions upon detection of pathogen risks to protect their offspring which lack the glands. We found an increase in the absolute amount of the benzoquinones (MBQ, EBQ) and their carrier 1-C15-ene in the secretions of wounded females 24 h after the treatment. The finding that particularly females may engage in niche construction fits theory, proposing more resource investment in immunity and longevity (Bateman 1948; Rolff 2002) or into brood protection (Trivers 1972) in females than in males. However, these effects lasted for less than 72 h after immune treatments. Potential metabolic costs associated with quinone production and individual survival costs caused by autotoxicity of quinones in high concentrations (Yezerski et al. 2004; Joop et al. 2014; Gokhale et al. 2017), may explain the transience of this effect. Moreover, on the group level enhanced quinone production may also increase herd-protection (Cotter and Kilner 2010), because under certain circumstances external immunity can be interpreted as public good (Yezerski et al. 2004; Joop et al. 2014; Gokhale et al. 2017). However, increased quinone levels in the environment may provide benefits especially to non-secreting *T. castaneum* conspecifics, which represents a social dilemma (Yezerski et al. 2004; Joop et al. 2014; Gokhale et al. 2017). Thus, there is likely a complex trade-off between benefits and various individual cost of quinone production, determining the time course of responses to immune experience. This study underlines the importance to distinguish between sexes and time points when exploring responses to immune challenge, potentially influencing niche construction.

Apart from serving as antimicrobials, benzoquinones and alkenes in stink gland secretions also lead to dose-dependent behavioral responses in conspecifics of *Tribolium* beetles (Suzuki 1985; Verheggen et al. 2007). For example, together with an aggregation pheromone, 4,8-dimethyldecanal, benzoquinones and alkenes mediate dispersal (Ogden 1969; Verheggen et al. 2007; Duehl et al. 2011). If wounding occurs at high population densities, an enhanced secretion of benzoquinones can signal overcrowding (Faustini and Burkholder 1987; Verheggen et al. 2007). In accordance with the results of our study, females appear particularly sensitive to perceive certain other cues, for example the secretions of other females in the flour, responded by upregulating their own quinone production (Sonleitner and Gutherie 1991; Khan et al. 2018). High amounts of quinones can even suppress the fecundity of conspecifics, which has huge consequences for population density regulation (Sonleitner and Gutherie 1991; Khan et al. 2018). We determined two important, yet unidentified compounds from secretion profiles using random forest analysis, which revealed significant sex-specific treatment effects on their relative amounts. The modulation of these compounds after immune challenge, potentially together with other cues, may serve yet unknown sex-specific functions, as described for pheromones in *T. molitor* (Rantala et al. 2002; Sadd et al. 2006; Nielsen and Holman 2012). Further quantification of quinone amounts actually present in the flour environment, together with identification of other potentially relevant compounds could help elucidating how the beetles regulate their environment and/or population density *via* their stink gland secretions.

Comparing different types of immune stimulation, in a previous study wounding alone (i.e., pricking control) resulted in weaker immune gene expression than treatment with alive *B. thuringiensis* bv. *tenebrionis* (Behrens et al. 2014). Here, however, the response of *T. castaneum* to priming was weaker than the response to wounding across both types of infochemicals, CHCs and quinones. Firstly, wounding may quickly establish a transient physiological state of heightened ‘alertness’, leading to responses with quinones as broadly-acting antimicrobials (Joop et al. 2014; Pedrini et al. 2015; Sawada et al. 2020) and potentially displayed to conspecifics in changed CHC profiles. Meanwhile, immune priming *via* injecting heat-killed bacteria may carry more information and induce additional physiological and immunological responses that mediate more tailored defenses against certain types of pathogens (Roth et al. 2009; Ferro et al. 2019). Secondly, potential competition for tyrosine precursors between both the quinone production and the phenoloxidase cascade activated upon bacterial challenge (Behrens et al. 2014; Joop et al. 2014; Otti et al. 2014), may result in a trade-off between external and internal immunity. Initially prioritizing individual internal immunity over external immunity, which confers protection also on adjacent conspecifics, may be beneficial for this non-social species. The aforementioned possible reasons together with the finding that 72 h after treatment primed beetles had the highest average amounts of quinones, hint to a delay or shift in immunological processes rather than absence of responses in primed beetles. This may indicate a possible more durable enhanced protection representing a ‘primed’ physiological state. Furthermore, the common patterns of a more pronounced response to wounding than to priming in both CHCs and quinone secretions may suggest that the modulation of both types of chemicals shares a common physiological cascade. For example, cytochrome P450 genes can be involved in CHC modification in insects (Chen et al. 2016) and have been shown to be crucial for quinone and alkene production in *T. castaneum* (Li et al. 2013; Lehmann 2015). Further analyses are needed to explore immune responses in CHCs and secretion compounds and their associated metabolic processes upon wounding *versus* priming over time.

In conclusion, our study highlights the potential role of chemical profiles in mediating the intertwined processes of immunity and niche construction in the red flour beetle. Immune stimulation triggered changes in potentially important infochemicals which can have fitness consequences for both the individual and the group. Furthermore, our results emphasize the high specificity of responses towards different types of immune stimuli and the importance to focus on sex-specific functions of chemicals. More generally, regarding the special lifestyle of a group-living but non-social insect, these results bear fundamental relevance for questions of common and specific mechanisms of disease control in groups and societies across taxa.

## Acknowledgements

We thank Robert Peuß for helpful comments on the manuscript.

## Data availability

Raw data and R codes are provided in the supplementary information. The GC-FID raw data will be released upon publication of the paper in the MetaboLights repository (Haug et al. 2013, 2020), being available with the accession number MTBLS2277 at www.ebi.ac.uk/metabolights/MTBLS2277.

## Contents of Online Supplementary Information

**Methods S1** *Bacteria Cultivation and Immunostimulant Preparation*

**Methods S2** *Details on Data Processing*

**Methods S3** *Details on Statistical Analyses*

**Fig. S1** NMDS plot and heatmaps of cuticular hydrocarbon profiles from naïve *T. castaneum*

**Fig. S2** NMDS plot of glandular secretion profiles from immune treated female *T. castaneum*

**Fig. S3** NMDS plot of glandular secretion profiles from immune treated male *T. castaneum*

**Fig. S4** Mean decrease in accuracy plot from random forest analysis of glandular secretion profiles of immune treated female *T. castaneum* and relative amounts of relevant feature RI 1015

**Fig. S5** Mean decrease in accuracy plot from random forest analysis of glandular secretion profiles of immune treated male *T. castaneum* and relative amounts of relevant feature RI 1473

**Table S1** Cuticular hydrocarbons and their relative composition in immune treated *T. castaneum*

**Table S2** Features putatively identified from glandular secretions of *T. castaneum* beetles

**Table S3** Candidate fatty acid synthase genes upregulated post wounding in *T. castaneum* larvae

## Statements and Declarations

### Author Contributions

JK and CM conceived the ideas. JK, CM, LJT, LKL and RR planned the experimental procedure. LKL and RR carried out the experiments and analyzed the raw data together with LJT. BM, LJT and LKL carried out the statistical analyses. All authors contributed to writing of the article and approved the submitted version.

## REFERENCES

Alabi T, Michaud JP, Arnaud L, Haubruge E (2008) A comparative study of cannibalism and predation in seven species of flour beetle. Ecol Entomol. https://doi.org/10.1111/j.1365-2311.2008.01020.x

Alciatore G, Ugelvig LV, Frank E, et al (2021) Immune challenges increase network centrality in a queenless ant. Proc Biol Sci 288:20211456. https://doi.org/10.1098/rspb.2021.1456

Alnajim I, Du X, Lee B, et al (2019) New method of analysis of lipids in *Tribolium castaneum* (Herbst) and *Rhyzopertha dominica* (Fabricius) insects by direct immersion solid-phase microextraction (DI-SPME) coupled with GC–MS. Insects 10:363

Altincicek B, Knorr E, Vilcinskas A (2008) Beetle immunity: Identification of immune-inducible genes from the model insect *Tribolium castaneum*. Dev Comp Immunol 32:585–595. https://doi.org/10.1016/j.dci.2007.09.005

Awater-Salendo S, Schulz H, Hilker M, Fürstenau B (2020) The importance of methyl-branched cuticular hydrocarbons for successful host recognition by the larval ectoparasitoid *Holepyris sylvanidis*. J Chem Ecol 46:1032–1046. https://doi.org/10.1007/s10886-020-01227-w

Baracchi D, Fadda A, Turillazzi S (2012) Evidence for antiseptic behaviour towards sick adult bees in honey bee colonies. J Insect Physiol 58:1589–1596. https://doi.org/10.1016/j.jinsphys.2012.09.014

Bateman AJ (1948) Intra-sexual selection in *Drosophila*. Heredity 2:349–368. https://doi.org/10.1038/hdy.1948.21

Behrens S, Peuß R, Milutinović B, et al (2014) Infection routes matter in population-specific responses of the red flour beetle to the entomopathogen *Bacillus thuringiensis*. BMC Genomics 15:445. https://doi.org/10.1186/1471-2164-15-445

Behringer V, Preis A, Wu DF, et al (2020) Urinary cortisol increases during a respiratory outbreak in wild chimpanzees. Front Vet Sci 7:485. https://doi.org/10.3389/fvets.2020.00485

Benjamini Y, Hochberg Y (1995) Controlling the false discovery rate: A practical and powerful approach to multiple testing. J R Stat Soc 57:289–300. https://doi.org/10.1111/j.2517-6161.1995.tb02031.x

Blomquist GJ, Ginzel MD (2021) Chemical ecology, biochemistry, and molecular biology of insect hydrocarbons. Annu Rev Entomol 66:45–60. https://doi.org/10.1146/annurev-ento-031620-071754

Blum MS (1981) 1,4-Quinones and hydroquinones. In: Chemical defenses of arthropods. Elsevier, pp 183–205

Campbell JF, Athanassiou CG, Hagstrum DW, Zhu KY (2022) *Tribolium castaneum*: A model insect for fundamental and applied research. Annu Rev Entomol 67:347–365. https://doi.org/10.1146/annurev-ento-080921-075157

Chapman RN (1926) Inhibiting the process of metamorphosis in the confused flour beetle (*Tribolium confusum*, Duval). J Exp Zool 45:293–299. https://doi.org/10.1002/jez.1400450110

Chen N, Fan Y-L, Bai Y, et al (2016) Cytochrome P450 gene, CYP4G51, modulates hydrocarbon production in the pea aphid, *Acyrthosiphon pisum*. Insect Biochem Mol Biol 76:84–94. https://doi.org/10.1016/j.ibmb.2016.07.006

Clutton-Brock TH (1991) The evolution of parental care. In: The Evolution of Parental Care. Princeton University Press.

Cotter SC, Kilner RM (2010) Personal immunity versus social immunity. Behav Ecol 21:663–668. https://doi.org/10.1093/beheco/arq070

Cremer S, Pull CD, Fürst MA (2018) Social immunity: Emergence and evolution of colony-level disease protection. Annu Rev Entomol 63:105–123. https://doi.org/10.1146/annurev-ento-020117-043110

Csata E, Timuş N, Witek M, et al (2017) Lock-picks: fungal infection facilitates the intrusion of strangers into ant colonies. Sci Rep 7:46323. https://doi.org/10.1038/srep46323

De Moraes CM, Stanczyk NM, Betz HS, et al (2014) Malaria-induced changes in host odors enhance mosquito attraction. Proc Natl Acad Sci U S A 111:11079–11084. https://doi.org/10.1073/pnas.1405617111

Duehl AJ, Arbogast RT, Teal PEA (2011) Density-related volatile emissions and responses in the red flour beetle, *Tribolium castaneum*. J Chem Ecol 37:525–532. https://doi.org/10.1007/s10886-011-9942-3

Faustini DL, Burkholder WE (1987) Quinone-aggregation pheromone interaction in the red flour beetle. Anim Behav 35:601–603. https://doi.org/10.1016/S0003-3472(87)80289-0

Ferro K, Ferro D, Corrà F, et al (2017) Cu,Zn Superoxide dismutase genes in: Evolution, molecular characterisation, and gene expression during immune priming. Front Immunol 8:1811. https://doi.org/10.3389/fimmu.2017.01811

Ferro K, Peuß R, Yang W, et al (2019) Experimental evolution of immunological specificity. Proc Natl Acad Sci U S A 116:20598–20604. https://doi.org/10.1073/pnas.1904828116

Gallagher JD, Siva-Jothy MT, Evison SEF (2018) Social cues trigger differential immune investment strategies in a non-social insect, *Tenebrio molitor*. Biology Letters 14:20170709

Gokhale CS, Traulsen A, Joop G (2017) Social dilemma in the external immune system of the red flour beetle? It is a matter of time. Ecol Evol 7:6758–6765. https://doi.org/10.1002/ece3.3198

Greenwood JM, Milutinović B, Peuß R, et al (2017) Oral immune priming with *Bacillus thuringiensis* induces a shift in the gene expression of Tribolium castaneum larvae. BMC Genomics 18:. https://doi.org/10.1186/s12864-017-3705-7

Haug K, Cochrane K, Nainala VC, et al (2020) MetaboLights: a resource evolving in response to the needs of its scientific community. Nucleic Acids Res 48:D440–D444. https://doi.org/10.1093/nar/gkz1019

Haug K, Salek RM, Conesa P, et al (2013) MetaboLights--an open-access general-purpose repository for metabolomics studies and associated meta-data. Nucleic Acids Res 41:D781–6. https://doi.org/10.1093/nar/gks1004

Hernández López J, Riessberger-Gallé U, Crailsheim K, Schuehly W (2017) Cuticular hydrocarbon cues of immune-challenged workers elicit immune activation in honeybee queens. Mol Ecol 26:3062–3073. https://doi.org/10.1111/mec.14086

Holze H, Schrader L, Buellesbach J (2021) Advances in deciphering the genetic basis of insect cuticular hydrocarbon biosynthesis and variation. Heredity 126:219–234. https://doi.org/10.1038/s41437-020-00380-y

Jallon J-M, David JR (1987) Variations in cuticular hydrocarbons among the eight species of the *Drosophila melanogaster* subgroup. Evolution 41:294–302. https://doi.org/10.1111/j.1558-5646.1987.tb05798.x

Joop G, Roth O, Schmid-Hempel P, Kurtz J (2014) Experimental evolution of external immune defences in the red flour beetle. J Evol Biol 27:1562–1571. https://doi.org/10.1111/jeb.12406

Kappeler PM, Cremer S, Nunn CL (2015) Sociality and health: impacts of sociality on disease susceptibility and transmission in animal and human societies. Philos Trans R Soc Lond B Biol Sci 370:. https://doi.org/10.1098/rstb.2014.0116

Khan I, Prakash A, Agashe D (2017) Experimental evolution of insect immune memory versus pathogen resistance. Proc Biol Sci 284:20171583. https://doi.org/10.1098/rspb.2017.1583

Khan I, Prakash A, Issar S, et al (2018) Female density-dependent chemical warfare underlies fitness effects of group sex ratio in flour beetles. Am Nat 191:306–317. https://doi.org/10.1086/695806

Kolde R (2019) pheatmap: Pretty Heatmaps. R package version 1.0.12

Kováts E (1958) Gas-chromatographische Charakterisierung organischer Verbindungen. Teil 1: Retentionsindices aliphatischer Halogenide, Alkohole, Aldehyde und Ketone. Helvetica Chimica Acta 41:1915–1932

Lehmann S (2015) Biology of odoriferous defensive stink glands of the red flour beetle *Tribolium castaneum*. Niedersächsische Staats-und Universitätsbibliothek Göttingen

Leonhardt SD, Menzel F, Nehring V, Schmitt T (2016) Ecology and evolution of communication in social insects. Cell 164:1277–1287. https://doi.org/10.1016/j.cell.2016.01.035

Li J, Lehmann S, Weißbecker B, et al (2013) Odoriferous Defensive stink gland transcriptome to identify novel genes necessary for quinone synthesis in the red flour beetle, *Tribolium castaneum*. PLoS Genet 9:e1003596. https://doi.org/10.1371/journal.pgen.1003596

Little TJ, Kraaijeveld AR (2004) Ecological and evolutionary implications of immunological priming in invertebrates. Trends Ecol Evol 19:58–60. https://doi.org/10.1016/j.tree.2003.11.011

Lockey KH (1978) Hydrocarbons of adult *Tribolium castaneum* hbst. and Tribolium confusum duv. (coleoptera: tenebrionidae). Comparative Biochemistry and Physiology Part B: Comparative Biochemistry 61:401–407

Loconti JD, Roth LM (1953) Composition of the odorous secretion of *Tribolium castaneum*. Annals of the Entomological Society of America 46:281–289

Martinez Arbizu P (2020) pairwiseAdonis: Pairwise multilevel comparison using adonis. R package version 0.4

Milutinović B, Kurtz J (2016) Immune memory in invertebrates. Semin Immunol 28:328–342. https://doi.org/10.1016/j.smim.2016.05.004

Milutinović B, Schmitt T (2022) Chemical cues in disease recognition and their immunomodulatory role in insects. Curr Opin Insect Sci 50:100884. https://doi.org/10.1016/j.cois.2022.100884

Milutinović B, Stolpe C, Peuβ R, et al (2013) The red flour beetle as a model for bacterial oral infections. PLoS One 8:e64638. https://doi.org/10.1371/journal.pone.0064638

Müller C, Caspers BA, Gadau J, Kaiser S (2020) The power of infochemicals in mediating individualized niches. Trends Ecol Evol 35:981–989. https://doi.org/10.1016/j.tree.2020.07.001

Müller T, Müller C (2016) Consequences of mating with siblings and nonsiblings on the reproductive success in a leaf beetle. Ecol Evol 6:3185–3197. https://doi.org/10.1002/ece3.2103

Nelson DR (1993) Methyl-branched lipids in insects. Insect lipids: chemistry, biochemistry and biology 271–315

Netea MG, Joosten LAB, Latz E, et al (2016) Trained immunity: A program of innate immune memory in health and disease. Science 352:aaf1098. https://doi.org/10.1126/science.aaf1098

Nielsen ML, Holman L (2012) Terminal investment in multiple sexual signals: immune-challenged males produce more attractive pheromones. Functional Ecology 26:20–28

Odling-Smee J, Erwin DH, Palkovacs EP, et al (2013) Niche construction theory: a practical guide for ecologists. Q Rev Biol 88:4–28. https://doi.org/10.1086/669266

Ogden JC (1969) Effect of Components of conditioned medium on behavior in *Tribolium confusum*. Physiol Zool 42:266–274. https://doi.org/10.1086/physzool.42.3.30155490

Oksanen J, Blanchet FG, Friendly M, et al (2020) vegan: Community Ecology Package. R package version 2.5-7. 2020

Ottensmann M, Stoffel MA, Nichols HJ, Hoffman JI (2018) GCalignR: An R package for aligning gas-chromatography data for ecological and evolutionary studies. PLoS One 13:e0198311. https://doi.org/10.1371/journal.pone.0198311

Otti O, Tragust S, Feldhaar H (2014) Unifying external and internal immune defences. Trends Ecol Evol 29:625–634. https://doi.org/10.1016/j.tree.2014.09.002

Park T, Mertz DB, Grodzinski W, Prus T (1965) Cannibalistic predation in populations of flour beetles. Physiol Zool 38:289–321. https://doi.org/10.1086/physzool.38.3.30152840

Pedrini N, Ortiz-Urquiza A, Huarte-Bonnet C, et al (2015) Tenebrionid secretions and a fungal benzoquinone oxidoreductase form competing components of an arms race between a host and pathogen. Proc Natl Acad Sci U S A 112:E3651–60. https://doi.org/10.1073/pnas.1504552112

Peterson R (2021) Finding optimal normalizing transformations via bestNormalize. R J 13:310. https://doi.org/10.32614/rj-2021-041

Peuß R, Eggert H, Armitage SAO, Kurtz J (2015) Downregulation of the evolutionary capacitor Hsp90 is mediated by social cues. Proc Biol Sci 282:. https://doi.org/10.1098/rspb.2015.2041

Prendeville HR, Stevens L (2002) Microbe inhibition by *Tribolium* flour beetles varies with beetle species, strain, sex, and microbe group. J Chem Ecol 28:1183–1190. https://doi.org/10.1023/a:1016281600915

Pull CD, Ugelvig LV, Wiesenhofer F, et al (2018) Destructive disinfection of infected brood prevents systemic disease spread in ant colonies. Elife 7:. https://doi.org/10.7554/eLife.32073

R Core Team (2020) R: A language and environment for statistical computing (Version 4.0.3, R Foundation for Statistical Computing, Vienna, Austria, 2020)

Rantala MJ, Jokinen I, Kortet R, et al (2002) Do pheromones reveal male immunocompetence? Proceedings of the Royal Society of London. Series B: Biological Sciences 269:1681–1685

Richard F-J, Aubert A, Grozinger CM (2008) Modulation of social interactions by immune stimulation in honey bee, *Apis mellifera*, workers. BMC Biol 6:50. https://doi.org/10.1186/1741-7007-6-50

Rolff J (2002) Bateman’s principle and immunity. Proc Biol Sci 269:867–872. https://doi.org/10.1098/rspb.2002.1959

Rosengaus RB, Maxmen AB, Coates LE, Traniello JFA (1998) Disease resistance: a benefit of sociality in the dampwood termite *Zootermopsis angusticollis* (Isoptera: Termopsidae). Behavioral Ecology and Sociobiology 44:125–134

Roth LM, Howland RB (1941) Studies on the gaseous secretion of *Tribolium confusum* Duval I. abnormalities produced in Tribolium confusum Duval by exposure to a secretion given off by the adults. Ann Entomol Soc Am 34:151–175

Roth O, Sadd BM, Schmid-Hempel P, Kurtz J (2009) Strain-specific priming of resistance in the red flour beetle, Tribolium castaneum. Proc Biol Sci 276:145–151. https://doi.org/10.1098/rspb.2008.1157

Rutherford SL, Lindquist S (1998) Hsp90 as a capacitor for morphological evolution. Nature 396:336–342. https://doi.org/10.1038/24550

Sadd B, Holman L, Armitage H, et al (2006) Modulation of sexual signalling by immune challenged male mealworm beetles (Tenebrio molitor, L.): evidence for terminal investment and dishonesty. Journal of Evolutionary Biology 19:321–325

Sawada M, Sano T, Hanakawa K, et al (2020) Benzoquinone synthesis-related genes of *Tribolium castaneum* confer the robust antifungal host defense to the adult beetles through the inhibition of conidial germination on the body surface. J Invertebr Pathol 169:107298. https://doi.org/10.1016/j.jip.2019.107298

Sokoloff A (1974) The biology of Tribolium: with special emphasis on genetic aspects. Oxford University Press, UK

Sonleitner FJ, Gutherie J (1991) Factors affecting oviposition rate in the flour beetle *Tribolium castaneum* and the origin of the population regulation mechanism. Res Popul Ecol 33:1–11. https://doi.org/10.1007/bf02514569

Stockmaier S, Stroeymeyt N, Shattuck EC, et al (2021) Infectious diseases and social distancing in nature. Science 371:. https://doi.org/10.1126/science.abc8881

Strobl C, Hothorn T, Zeileis A (2009a) Party on! A new, conditional variable importance measure for random forests Available in the party Package. R J 1:14–17

Strobl C, Malley J, Tutz G (2009b) An introduction to recursive partitioning: Rationale, application, and characteristics of classification and regression trees, bagging, and random forests. Psychol Methods 14:323–348. https://doi.org/10.1037/a0016973

Suzuki T (1985) Presence of another aggregation substance(s) in the frass of the red flour beetles, *Tribolium castaneum* (Coleoptera: Tenebrionidae). Appl Entomol Zool 20:90–91. https://doi.org/10.1303/aez.20.90

Swanson JAI, Torto B, Kells SA, et al (2009) Odorants that induce hygienic behavior in honeybees: identification of volatile compounds in chalkbrood-infected honeybee larvae. J Chem Ecol 35:1108–1116. https://doi.org/10.1007/s10886-009-9683-8

Tate AT, Graham AL (2017) Dissecting the contributions of time and microbe density to variation in immune gene expression. Proc Biol Sci 284:. https://doi.org/10.1098/rspb.2017.0727

Thomas ML, Simmons LW (2008) Sexual dimorphism in cuticular hydrocarbons of the Australian field cricket *Teleogryllus oceanicus* (Orthoptera: Gryllidae). J Insect Physiol 54:1081–1089. https://doi.org/10.1016/j.jinsphys.2008.04.012

Trivers R (1972) Parental investment and sexual selection. Sexual Selection & the Descent of Man, Aldine de Gruyter, New York 136–179

Van Wyk JH, Hodson AC, Christensen CM (1959) Microflora associated with the confused flour beetle, *Tribolium Confusum1*. Annals of the Entomological Society of America 52:452–463

van Zweden JS, d’Ettorre P (2010) Nestmate recognition in social insects and the role of hydrocarbons. Insect hydrocarbons: biology, biochemistry and chemical ecology 11:222–243

Verheggen F, Ryne C, Olsson POC, et al (2007) Electrophysiological and behavioral activity of secondary metabolites in the confused flour beetle, *Tribolium confusum*. J Chem Ecol 33:525–539. https://doi.org/10.1007/s10886-006-9236-3

Via S (1999) Cannibalism facilitates the use of a novel environment in the flour beetle, *Tribolium castaneum*. Heredity 82 (Pt 3):267–275. https://doi.org/10.1038/sj.hdy.6884820

Villaverde ML, Luciana Villaverde M, Patricia Juárez M, Mijailovsky S (2007) Detection of *Tribolium castaneum* (Herbst) volatile defensive secretions by solid phase microextraction– capillary gas chromatography (SPME-CGC). Journal of Stored Products Research 43:540–545

Wang L, Elliott B, Jin X, et al (2015) Antimicrobial properties of nest volatiles in red imported fire ants, *Solenopsis invicta* (Hymenoptera: Formicidae). Sci Nat 102:66. https://doi.org/10.1007/s00114-015-1316-1

Wegner KM, Berenos C, Schmid-Hempel P (2008) Nonadditive genetic components in resistance of the red flour beetle *Tribolium castanaeum* against parasite infection. Evolution 62:2381–2392. https://doi.org/10.1111/j.1558-5646.2008.00444.x

Yew JY, Chung H (2015) Insect pheromones: An overview of function, form, and discovery. Prog Lipid Res 59:88–105. https://doi.org/10.1016/j.plipres.2015.06.001

Yezerski A, Ciccone C, Rozitski J, Volingavage B (2007) The effects of a naturally produced benzoquinone on microbes common to flour. Journal of Chemical Ecology 33:1217–1225

Yezerski A, Gilmor TP, Stevens L (2000) Variation in the production and distribution of substituted benzoquinone compounds among genetic strains of the confused flour beetle, *Tribolium confusum*. Physiol Biochem Zool 73:192–199. https://doi.org/10.1086/316733

Yezerski A, Gilmor TP, Stevens L (2004) Genetic analysis of benzoquinone production in *Tribolium confusum*. J Chem Ecol 30:1035–1044. https://doi.org/10.1023/b:joec.0000028465.37658.ae

