## Supplements for "Immune Stimulation *via* Wounding Alters Chemical Profiles of Adult *Tribolium castaneum*"

for the manuscript

### Contents

**Methods S1** *Bacteria Cultivation and Immunostimulant Preparation*

**Methods S2** *Details on Data Processing*

**Methods S3** *Details on Statistical Analyses*

**Fig. S1** NMDS plot and heatmaps of cuticular hydrocarbon profiles from naïve *T. castaneum*

**Fig. S2** NMDS plot of glandular secretion profiles from immune treated female *T. castaneum*

**Fig. S3** NMDS plot of glandular secretion profiles from immune treated male *T. castaneum*

**Fig. S4** Mean decrease in accuracy plot from random forest analysis of glandular secretion profiles of immune treated female *T. castaneum* and relative amounts of relevant feature RI 1015

**Fig. S5** Mean decrease in accuracy plot from random forest analysis of glandular secretion profiles of immune treated male *T. castaneum* and relative amounts of relevant feature RI 1473

**Table S1** Cuticular hydrocarbons and their relative composition in immune treated *T. castaneum*

**Table S2** Features putatively identified from glandular secretions of *T. castaneum* beetles

**Table S3** Candidate fatty acid synthase genes upregulated post wounding in *T. castaneum* larvae

**Methods S1**

*Bacteria Cultivation and Immunostimulant Preparation*. To prepare the bacterial culture, 100 µL of a *Bacillus thuringiensis* bv. *tenebrionis* freezer stock (stored at -80 ºC) was inoculated into a 500 mL baffled flask containing 50 mL Luria Broth (Carl Roth GmbH + Co. KG) and the culture was incubated at 30 ºC, 200 rpm for 15 h in dark. Subsequently, the culture was transferred into a 50 mL Falcon conical tube and centrifuged at 3,645 × *g* at 4 ºC for 10 min. The bacterial pellet was washed once with phosphate buffered saline (PBS, Calbiochem®) and the concentration adjusted to $1 \times 10^9$ cells $mL^{-1}$. PBS (as injection/wounding control) and bacteria solution (for heat-killed *B. thuringiensis* bv. *tenebrionis* injecting treatment, 'priming') were heated at 95 ºC for 30 min to heat-kill the bacteria. All procedures were carried out under a horizontal laminar air-flow hood to keep the conditions sterile.

**Methods S2**

*Details on Data Processing.* Data pre-processing was done using the GCsolution Postrun software (Version 2.30.00, Shimadzu), exporting peak areas with at least 100 (CHCs) and 5 counts (stink gland secretion), respectively. Retention time alignment of peaks was done in R (version 4.0.3, (R Core Team 2020) using the package *GCalignR* (Ottensmann et al. 2018), allowing a maximum shift in retention time by 0.05 min. Features that could not be distinguished from those occurring also in blank samples were excluded. The resulting datasets were both normalised to the internal standard and only features detected in at least half of the replicates per treatment and sex (and time point in case of stink gland secretions) were considered for further analysis. When relative values were used for analysis, these were calculated by expressing the amount of a certain feature as the proportion of the total amount of features considered for a sample.

**Methods S3**

*Details on Statistical Analyses.* All statistical analyses were done using R (version 4.0.3). To visualise dissimilarities between groups of beetles, non-metric multidimensional scaling (NMDS) was applied to all datasets using the *metaMDS* function in the package *vegan* (version 2.5-7, (Oksanen et al. 2020). For this, datasets were transformed using the Wisconsin double standardisation (package *vegan*) of the square root values and pairwise dissimilarities were calculated based on the Kulczynski index. NMDS analysis was performed on relative feature abundances, separately for females and males, and separately for the two time points in the analysis of gland secretions. For the analysis of CHC profiles, naïve males and females were additionally compared in a separate analysis to visualise CHC profile differences attributed to sex. Permutational multivariate analysis of variance (PERMANOVA) was performed using the *adonis* function (package *vegan*) to test for significance of the on the dissimilarities displayed by NMDS plots. The same transformations and dissimilarity indices as for NMDS were employed and 10,000 permutations were used. The *betadisper* and *permutest* (10,000 permutations) functions (package *vegan*) were used to confirm the homogeneity of the multivariate variance spread among the treatment groups. If significant treatment effects were found in the PERMANOVA analyses of females and males, pairwise comparisons between treatment groups were performed (package *pairwiseAdonis*, (Martinez Arbizu 2020) with correction of *P*-values using the Benjamini-Hochberg ('BH', (Benjamini and Hochberg 1995) procedure (a = 0.05).

For the secretions, we further used the datasets acquired 24 h after treatment and identified features that differed between treatments in both females and males using a conditional random forest classification (seed 245, n trees = 500, n variables per split = 5) (Strobl et al. 2009a, b; De Moraes et al. 2014). The analysis was done on relative feature amounts to control for different total amounts of secretions due to individual variations and random gland emptying before and during experimental procedures, thereby capturing changes in the proportion of features to one another. Random forest analysis identified one feature as important for each sex, whose relative concentrations were further analysed in post hoc testing. We tested for data normality, heteroscedasticity, presence of multicollinearity and influential cases and obtained diagnostics plots (Q-Q plots and histograms of model residuals distribution and a plot to estimate non-linear patterns of residuals). Not meeting the model assumptions, the two features were analysed with non-parametric Kruskal-Wallis tests and equivalent pairwise comparisons between treatment groups. We further analysed the effect of treatments on the absolute concentration of secretion compounds with known biological activity, EBQ, MBQ and 1-pentadecene (1-C15-ene) separately for both sexes at both time points using linear models (LMs). We tested the model assumptions as described above and employed the package *bestNormalize* (Peterson 2021) to determine the appropriate transformation methods to obtain normality. The following transformations were used: MBQ, 24 h: sqrt(x + a), 72 h: asinh(x); EBQ, 24 h: Box Cox, 72 h: orderNorm; 1-C15-ene, 24 h and 72 h: Box Cox. When significant treatment effects were founds, Tukey HSD post hoc comparisons were performed between treatment groups with *P*-value corrections using the BH procedure (a = 0.05).

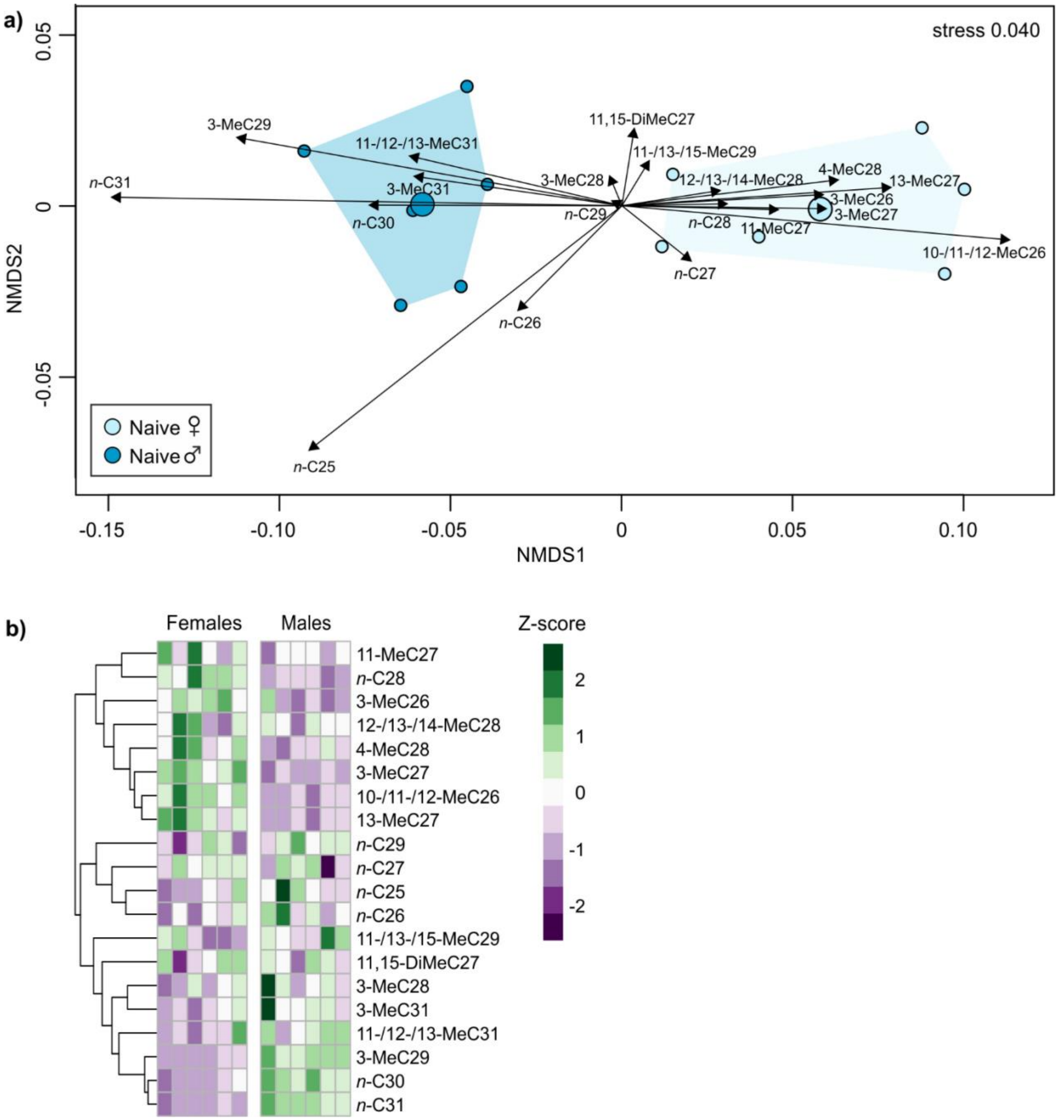


**Fig. S1** Non-metric multidimensional scaling plot (**a**) and heatmaps (**b**) of cuticular hydrocarbon (CHC) profiles from naïve female and male *Tribolium castaneum* beetles (n = 6 per sex). All plots are based on relative amounts of 20 individual CHCs. For NMDS plots, datasets were transformed with Wisconsin square root double standardisation and dissimilarities are based on pairwise Kulczynki distances. Data points represent profiles of individual beetles, larger symbols show centroids for treatment groups. In (**a**) arrows range from zero to positions of expanded weighted average scores of CHCs. Heatmaps show differences in the CHC profiles of females versus males based on z-scores of individual CHCs. In heatmaps, clustering of CHCs is based on Euclidean distances.

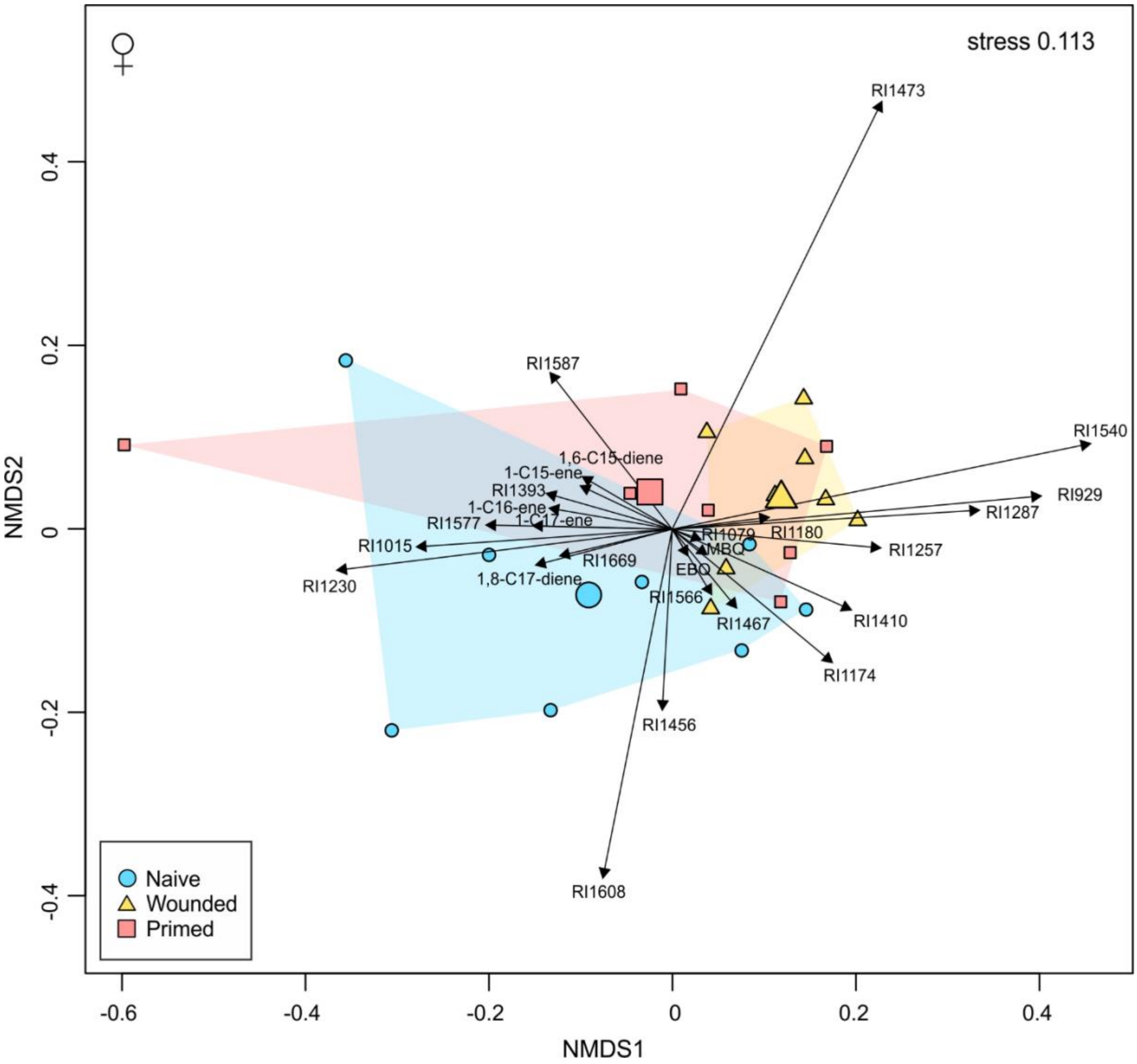


**Fig. S2** Non-metric multidimensional scaling plot of glandular secretion profiles from female *Tribolium castaneum* beetles of different immune treatment groups (n = 7-8 per treatment). Beetles were untreated ('Naïve'), or treated by injection of phosphate buffered saline ('Wounded') or of phosphate buffered saline with heat-killed *Bacillus thuringiensis* bv. *tenebrionis* ('Primed'), 24 h before extraction. Dataset comprised relative amounts of 26 chemical features (absolute amounts divided by the total amount of all features considered for a sample) and was transformed with Wisconsin square root double standardisation. Dissimilarities are based on pairwise Kulczynki distances. Labels are retention index (RI) values or the compound name; ethyl-1,4-benzoquinone (EBQ), methyl-1,4-benzoquinone (MBQ). Data points represent profiles of individual beetles, larger symbols show centroids for treatment groups. Arrows range from zero to positions of expanded weighted average scores of features. The stress value is given at the top right.

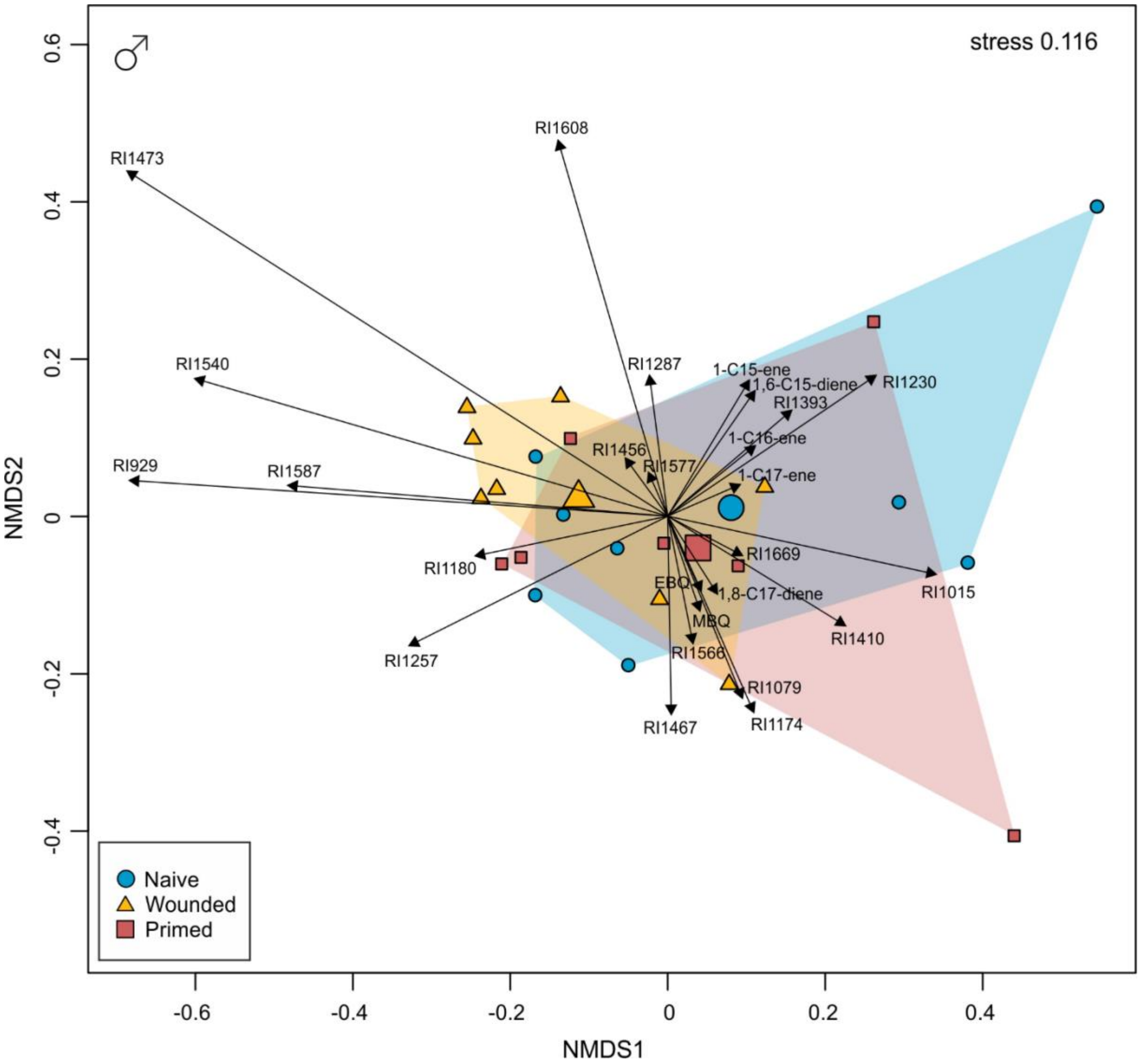


**Fig. S3** Non-metric multidimensional scaling plot of glandular secretion profiles from male *Tribolium castaneum* beetles of different immune treatment groups (n = 7-8 per treatment). Beetles were untreated ('Naïve'), or treated by injection of phosphate buffered saline ('Wounded') or of phosphate buffered saline with heat-killed *Bacillus thuringiensis* bv. *tenebrionis* ('Primed'), 24 h before extraction. Dataset comprised relative amounts of 26 chemical features (absolute amounts divided by the total amount of all features considered for a sample) and was transformed with Wisconsin square root double standardisation. Dissimilarities are based on pairwise Kulczynki distances. Labels are retention index (RI) values or the compound name; ethyl-1,4-benzoquinone (EBQ), methyl-1,4-benzoquinone (MBQ). Data points represent profiles of individual beetles, larger symbols show centroids for treatment groups. Arrows range from zero to positions of expanded weighted average scores of features. The stress value is given at the top right.

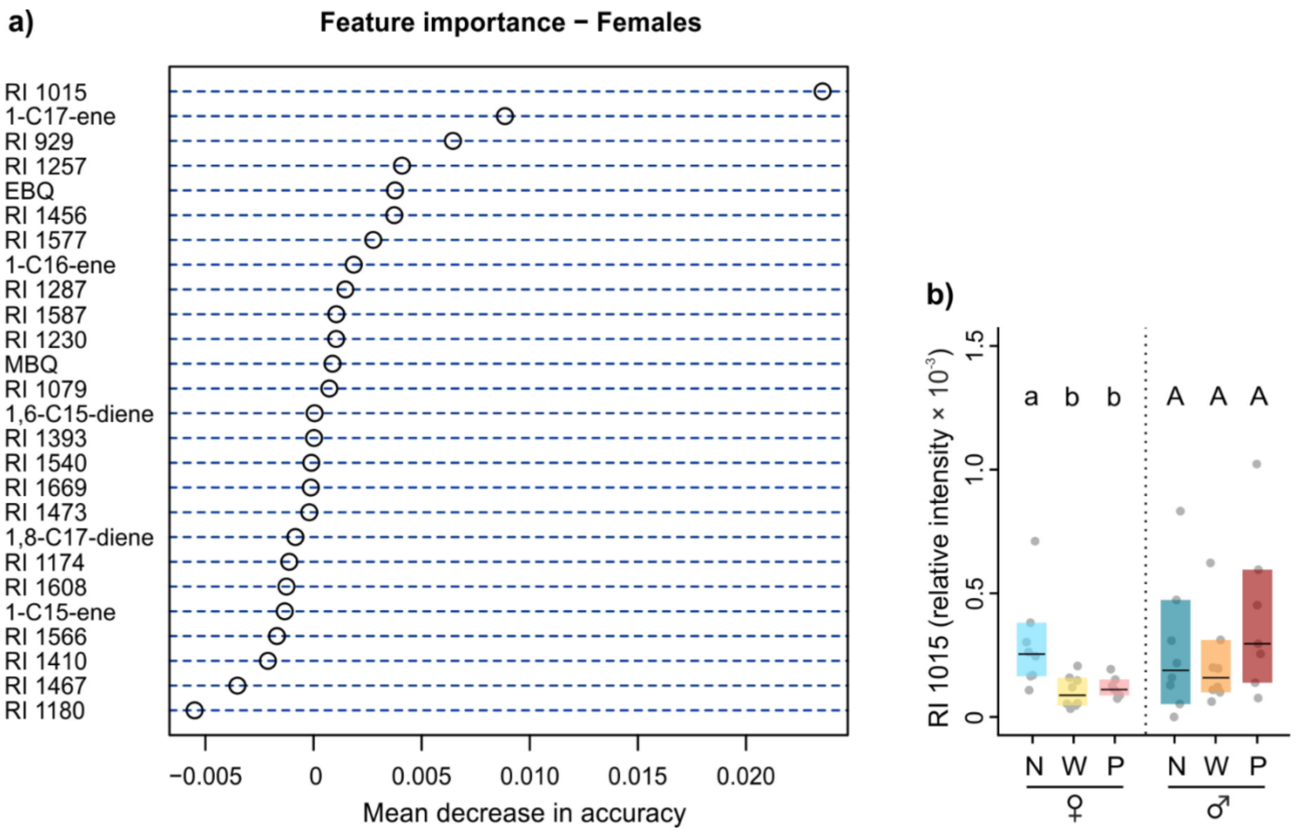


**Fig. S4 a**) Mean decrease in accuracy plot from conditional random forest (RF) analysis of features from glandular secretions of female *Tribolium castaneum* beetles of different immune treatment groups (n = 7-8 per treatment). Beetles were untreated (naïve, 'N'), or treated by injection of phosphate buffered saline (wounded, 'W') or of phosphate buffered saline with heat-killed *Bacillus thuringiensis* bv. *tenebrionis* (primed, 'P'), 24 h before extraction. Relative amounts, obtained by dividing absolute amounts by the total amount of all features considered for a sample were used for analysis. Labels are retention index (RI) values or the compound name; ethyl-1,4-benzoquinone (EBQ), methyl-1,4-benzoquinone (MBQ). **b**) Relative amounts of the feature with RI 1015, suggested as most important for separation of treatment groups within females by RF analysis. Dots represent values of individual beetles, boxes are confidence intervals, lines are medians. Different letters above boxes show significant differences ($P$ < 0.05) between treatment groups within females (lower case letters) and males (upper case letters) in *Kruskal-Wallis tests* after pairwise comparisons.

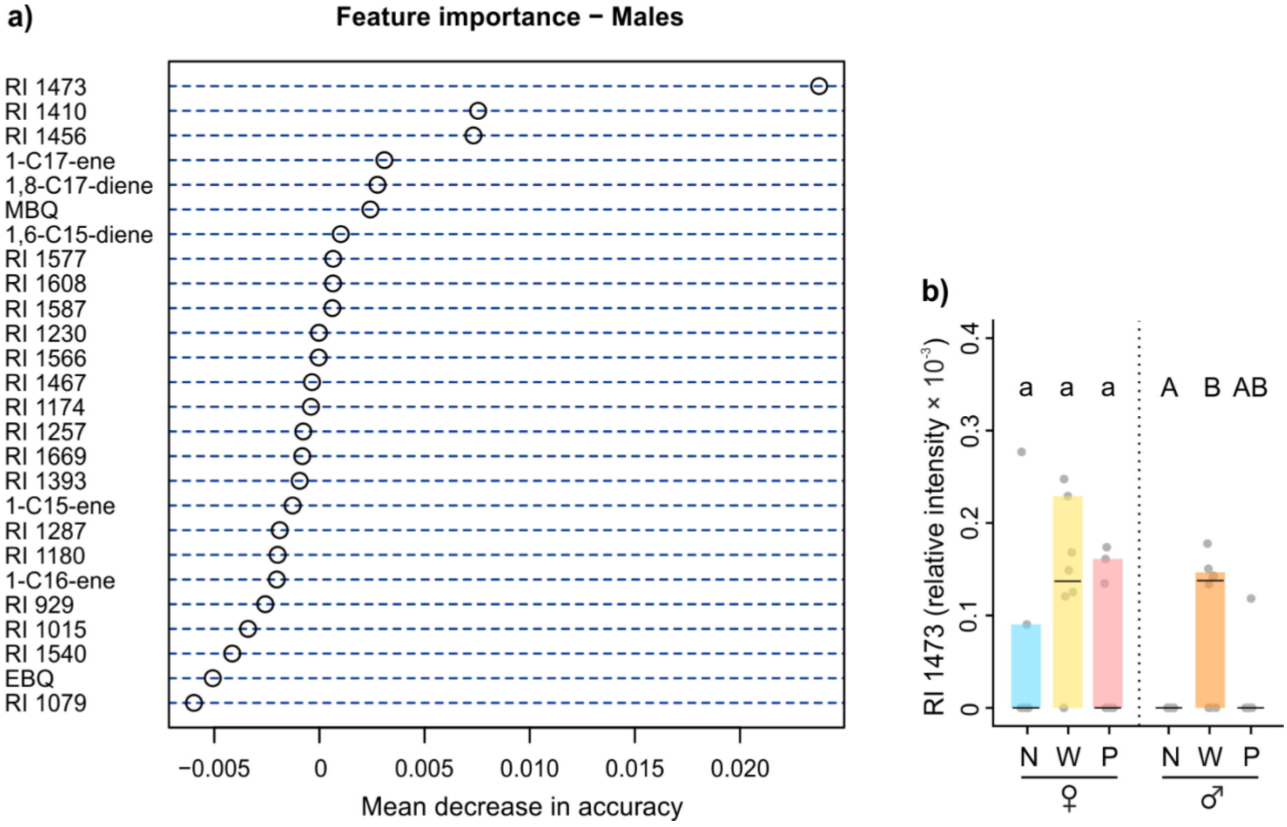


**Fig. S5 a**) Mean decrease in accuracy plot from conditional random forest (RF) analysis of features from glandular secretions of male *Tribolium castaneum* beetles of different immune treatment groups (n = 7-8 per treatment). Beetles were untreated (naïve, 'N'), or treated by injection of phosphate buffered saline (wounded, 'W') or of phosphate buffered saline with heat-killed *Bacillus thuringiensis* bv. *tenebrionis* (primed, 'P'), 24 h before extraction. Relative amounts, obtained by dividing absolute amounts by the total amount of all features considered for a sample were used for analysis. Labels are retention index (RI) values or the compound name; ethyl-1,4-benzoquinone (EBQ), methyl-1,4-benzoquinone (MBQ). **b**) Relative amounts of the feature with RI 1473, suggested as most important for separation of treatment groups within males by RF analysis. Dots represent values of individual beetles, boxes are confidence intervals, lines are medians. Different letters above boxes show significant differences ($P < 0.05$) between treatment groups within females (lower case letters) and males (upper case letters) in *Kruskal-Wallis tests* after pairwise comparisons.

**Table S1** Cuticular hydrocarbons (CHCs) and their relative composition in *Tribolium castaneum* beetles of different immune treatment groups (n = 6 per sex and treatment). Beetles were untreated ('Naïve'), or treated by injection of phosphate buffered saline ('Wounded') or of phosphate buffered saline with heat-killed *Bacillus thuringiensis* bv. *tenebrionis* ('Primed'), 18 h before extraction. Features were putatively identified by comparing their retention index (RI) value to (Lockey 1978), (Alnajim et al. 2019) and (Awater-Salendo et al. 2020). Forward slash between positions of methyl groups in CHC names indicates that positions are unclear or mixtures could not be separated. If literature suggested different identities, names were given based on better matching RI value (No. 12), better peak separation (No. 6, 7), or most recent publication (No. 3, 8, 11, 12, 15). For No. 19, potential positions of methyl groups are derived from suggestions in the text. The measured RIs of CHCs differed not more than ± 4 from the RIs proposed by literature.

| | | | Naive (% Mean ± SD) | | Wounded (% Mean ± SD) | | Primed (% Mean ± SD) | |
|---|---|---|---|---|---|---|---|---|
| No | RI | Name | Females | Males | Females | Males | Females | Males |
| 1 | 2498 | *n*-C25 | 0.4 ± 0.2 | 0.7 ± 0.3 | 0.4 ± 0.1 | 0.8 ± 0.1 | 0.4 ± 0.2 | 0.6 ± 0.2 |
| 2 | 2598 | *n*-C26 | 0.6 ± 0.1 | 0.8 ± 0.2 | 0.6 ± 0.1 | 0.9 ± 0.1 | 0.7 ± 0.2 | 0.8 ± 0.1 |
| 3 | 2629 | 10-/11-/12-MeC26 | 0.9 ± 0.2 | 0.5 ± 0.1 | 1.2 ± 0.2 | 0.7 ± 0.1 | 1.1 ± 0.2 | 0.7 ± 0.1 |
| 4 | 2669 | 3-MeC26 | 0.7 ± 0.1 | 0.5 ± 0.1 | 0.9 ± 0.1 | 0.7 ± 0.1 | 0.8 ± 0.1 | 0.6 ± 0.1 |
| 5 | 2698 | *n*-C27 | 10.2 ± 0.5 | 10.1 ± 1.3 | 9.7 ± 1.6 | 9.9 ± 0.9 | 10.1 ± 1.5 | 9.7 ± 1.2 |
| 6 | 2733 | 13-MeC27 | 20.0 ± 3.0 | 15.0 ± 1.5 | 26.3 ± 0.9 | 18.2 ± 1.4 | 24.7 ± 2.7 | 17.7 ± 1.3 |
| 7 | 2736 | 11-MeC27 | 0.9 ± 0.2 | 0.8 ± 0.1 | 1.2 ± 0.1 | 1.0 ± 0.1 | 1.0 ± 0.1 | 1.0 ± 0.1 |
| 8 | 2763 | 11,15-DiMeC27 | 5.5 ± 1.0 | 5.6 ± 0.8 | 5.3 ± 1.2 | 5.9 ± 0.8 | 5.2 ± 1.2 | 6.2 ± 1.4 |
| 9 | 2771 | 3-MeC27 | 13.8 ± 1.0 | 10.8 ± 0.6 | 14.0 ± 0.5 | 10.9 ± 1.2 | 13.9 ± 0.8 | 10.5 ± 0.3 |
| 10 | 2799 | *n*-C28 | 6.8 ± 0.3 | 6.4 ± 0.2 | 6.6 ± 0.2 | 6.5 ± 0.3 | 6.7 ± 0.5 | 6.4 ± 0.2 |
| 11 | 2827 | 12-/13-/14-MeC28 | 2.8 ± 0.4 | 2.7 ± 0.2 | 3.4 ± 0.2 | 3.3 ± 0.4 | 3.3 ± 0.2 | 3.3 ± 0.8 |
| 12 | 2853 | 4-MeC28 | 0.9 ± 0.2 | 0.7 ± 0.1 | 1.0 ± 0.1 | 0.9 ± 0.1 | 1.1 ± 0.2 | 0.7 ± 0.1 |
| 13 | 2870 | 3-MeC28 | 0.5 ± 0.1 | 0.6 ± 0.1 | 0.6 ± 0.1 | 0.6 ± 0.0 | 0.5 ± 0.1 | 0.6 ± 0.1 |
| 14 | 2898 | *n*-C29 | 21.2 ± 2.9 | 24.1 ± 1.7 | 18.0 ± 1.1 | 20.5 ± 2.2 | 19.5 ± 1.9 | 21.7 ± 2.2 |
| 15 | 2926 | 11-/13-/15-MeC29 | 3.2 ± 0.4 | 3.6 ± 0.4 | 4.0 ± 0.3 | 4.1 ± 0.3 | 3.9 ± 0.4 | 3.8 ± 0.5 |
| 16 | 2971 | 3-MeC29 | 4.0 ± 0.7 | 8.6 ± 1.2 | 3.5 ± 1.0 | 8.2 ± 0.6 | 3.6 ± 0.6 | 8.2 ± 0.9 |
| 17 | 2998 | *n*-C30 | 1.1 ± 0.2 | 1.8 ± 0.2 | 0.9 ± 0.1 | 1.7 ± 0.2 | 1.0 ± 0.1 | 1.7 ± 0.1 |
| 18 | 3098 | *n*-C31 | 1.9 ± 0.6 | 5.0 ± 0.7 | 1.5 ± 0.2 | 3.9 ± 0.4 | 1.5 ± 0.1 | 4.4 ± 0.5 |
| 19 | 3129 | 11-/12-/13-MeC31 | 0.7 ± 0.3 | 1.0 ± 0.2 | 0.6 ± 0.1 | 0.9 ± 0.1 | 0.6 ± 0.2 | 1.0 ± 0.1 |
| 20 | 3174 | 3-MeC31 | 0.4 ± 0.1 | 0.5 ± 0.2 | 0.3 ± 0.0 | 0.4 ± 0.1 | 0.3 ± 0.1 | 0.5 ± 0.1 |

**Table S2** Features putatively identified from glandular secretions of *Tribolium castaneum* beetles by comparing their retention index (RI) value to Li et al. 2013 and Lehmann 2015. No. 13 and 23 could not be detected in the dataset acquired 72 h after the treatment. Average proportions of all features are shown as percentage of the total amount of all features considered for a sample. The average proportions were calculated across females and males of all treatments and time points.

| No | RI | Name | Proportion [%] |
|---|---|---|---|
| 1 | 929 | Unidentified | <0.1 |
| 2 | 1015 | Unidentified | <0.1 |
| 3 | 1026 | Methyl-1,4-benzoquinone (MBQ) | 17.6 |
| 4 | 1079 | Unidentified | 0.1 |
| 5 | 1118 | Ethyl-1,4-benzoquinone (EBQ) | 22.2 |
| 6 | 1174 | Unidentified | 0.1 |
| 7 | 1180 | Unidentified | <0.1 |
| 8 | 1230 | Unidentified | <0.1 |
| 9 | 1257 | Unidentified | 0.1 |
| 10 | 1287 | Unidentified | <0.1 |
| 11 | 1393 | Unidentified | 0.1 |
| 12 | 1410 | Unidentified | <0.1 |
| 13 | 1456 | Unidentified | <0.1 |
| 14 | 1467 | Unidentified | 0.1 |
| 15 | 1473 | Unidentified | <0.1 |
| 16 | 1479 | 1,6-C15-diene | 1.8 |
| 17 | 1494 | 1-C15-ene | 42.4 |
| 18 | 1540 | Unidentified | <0.1 |
| 19 | 1566 | Unidentified | 1.5 |
| 20 | 1577 | Unidentified | 0.1 |
| 21 | 1587 | Unidentified | <0.1 |
| 22 | 1594 | 1-C16-ene | 0.3 |
| 23 | 1608 | Unidentified | <0.1 |
| 24 | 1669 | Unidentified | 0.4 |
| 25 | 1675 | 1,8-C17-diene | 8.5 |
| 26 | 1694 | 1-C17-ene | 4.7 |

**Table S3** Candidate fatty acid synthase genes upregulated 6 h and 18 h post wounding in *Tribolium castaneum* larvae (N: naïve, W: wounded (n = 32 × 3). FPKM values: Fragments per kilobase of exon per million fragments mapped.

| | | | | | 6 h | | | | 18 h | | | |
|---|---|---|---|---|---|---|---|---|---|---|---|---|
| | | | | | FPKM value | | | | FPKM value | | | |
| No | TC ID | InterPro description | Blast2GO description | Locus | N | W | $\log_2$ fold change | q value | N | W | $\log_2$ fold change | q value |
| 1 | TC000238 | Acyl transferase domain | fatty acid synthase-like | ChLG2: 18356249-18367184 | 51.8 | 74.6 | 0.5 | <0.01 | 60.7 | 80.2 | 0.4 | 0.04 |
| 2 | TC015399 | Acyl carrier protein-like | fatty acid synthase s-acetyltransferase | ChLG6: 351095-364842 | 4.1 | 7.6 | 0.9 | <0.01 | 6.0 | 16.0 | 1.4 | <0.01 |
| 3 | TC015400 | Acyl carrier protein-like | fatty acid synthase s-acetyltransferase | ChLG6: 367355-376905 | 6.7 | 19.9 | 1.6 | <0.01 | 13.8 | 36.4 | 1.4 | <0.01 |
| 4 | TC015337 | Acyl carrier protein-like | fatty acid synthase s-acetyltransferase | ChLG6: 241414-251114 | 8.0 | 14.2 | 0.8 | <0.01 | 13.9 | 31.0 | 1.2 | <0.01 |

*Note*. Adapted from “Infection routes matter in population-specific responses of the red flour beetle to the entomopathogen *Bacillus thuringiensis*” by Behrens S, Peuß R, Milutinović B, et al, 2014, BMC Genomics, 15, p. 445 (https://doi.org/10.1186/1471-2164-15-445). Copyright 2014 by Springer Nature

### References


Alnajim I, Du X, Lee B, et al (2019) New method of analysis of lipids in *Tribolium castaneum* (Herbst) and *Rhyzopertha dominic*a (Fabricius) insects by direct immersion solid-phase microextraction (DI-SPME) coupled with GC–MS. Insects 10:363

Awater-Salendo S, Schulz H, Hilker M, Fürstenau B (2020) The importance of methyl-branched cuticular hydrocarbons for successful host recognition by the larval ectoparasitoid *Holepyris sylvanidis*. J Chem Ecol 46:1032–1046. https://doi.org/10.1007/s10886-020-01227-w

Benjamini Y, Hochberg Y (1995) Controlling the false discovery rate: A practical and powerful approach to multiple testing. J R Stat Soc 57:289–300. https://doi.org/10.1111/j.2517-6161.1995.tb02031.x

De Moraes CM, Stanczyk NM, Betz HS, et al (2014) Malaria-induced changes in host odors enhance mosquito attraction. Proc Natl Acad Sci U S A 111:11079–11084. https://doi.org/10.1073/pnas.1405617111

Lehmann S (2015) Biology of odoriferous defensive stink glands of the red flour beetle *Tribolium castaneum*. Niedersächsische Staats-und Universitätsbibliothek Göttingen

Li J, Lehmann S, Weißbecker B, et al (2013) Odoriferous defensive stink gland transcriptome to identify novel genes necessary for quinone synthesis in the red flour beetle, *Tribolium castaneum*. PLoS Genet 9:e1003596. https://doi.org/10.1371/journal.pgen.1003596

Lockey KH (1978) Hydrocarbons of adult *Tribolium castaneum* hbst. and *Tribolium confusum* duv. (coleoptera: tenebrionidae). Comparative Biochemistry and Physiology Part B: Comparative Biochemistry 61:401–407

Martinez Arbizu P (2020) pairwiseAdonis: Pairwise multilevel comparison using adonis. R package version 0.4

Oksanen J, Blanchet FG, Friendly M, et al (2020) vegan: Community Ecology Package. R package version 2.5-7. 2020

Ottensmann M, Stoffel MA, Nichols HJ, Hoffman JI (2018) GCalignR: An R package for aligning gas-chromatography data for ecological and evolutionary studies. PLoS One 13:e0198311. https://doi.org/10.1371/journal.pone.0198311

Peterson R (2021) Finding optimal normalizing transformations via bestNormalize. R J 13:310. https://doi.org/10.32614/rj-2021-041

R Core Team (2020) R: A Language and Environment for Statistical Computing (Version 4.0.3, R Foundation for Statistical Computing, Vienna, Austria, 2020)

Strobl C, Hothorn T, Zeileis A (2009a) Party on! A New, Conditional Variable Importance Measure for Random Forests Available in the party Package. R J 1:14–17

Strobl C, Malley J, Tutz G (2009b) An introduction to recursive partitioning: Rationale, application, and characteristics of classification and regression trees, bagging, and random forests. Psychol Methods 14:323–348. https://doi.org/10.1037/a0016973
